# Computational modeling of breast cancer metastases predicts response to dual checkpoint inhibition and reveals spatially-informed treatments

**DOI:** 10.64898/2026.10.16.759478

**Authors:** Tyler King, Edgar Gonzalez, Evanthia T. Roussos Torres, Stacey D. Finley

## Abstract

In advanced breast cancers, the highly suppressive tumor microenvironment contributes to poor responses to immune checkpoint inhibition. The degree of suppression varies by site of disease, which can affect metastatic progression. To investigate this phenomenon, we have developed an agent-based model of cell-cell interactions within the tumor microenvironment, specifically tuned to *in vivo* data from murine breast-to-lung metastases. The model includes nine functional cell types (cancer cells, M0-, M1-, and M2-like macrophages, CD4+ T cells, T regulatory cells [Tregs], CD8+ T cells, NK cells, and MDSC-like cells). We incorporated a novel continuum model to describe exhaustion of CD8+ T and NK cells, where their exhaustion is impacted by cell-cell interactions in the TME via direct and indirect interactions. We also modeled the effect of the immune checkpoint inhibitors (ICI) anti-PD-1 and anti-CTLA-4, termed “dual ICIs”. We used this model to simulate the spatiotemporal trajectories of 19 metastases without treatment and with dual ICIs. The model predicted that dual ICIs lead to statistically significant reductions in cancer cell proliferation in seven of the metastases (37%), and spatial analyses of these metastases indicate they have structural similarities at multiple scales of tumor organization. Additionally, spatial analysis suggests that high colocalization of CD8+ T cells with Tregs could prevent dual ICIs from being effective, and simulation of Treg blockade combined with dual ICIs led to 15 total metastases having a statistically significant treatment effect (80% total). Thus, our model provides a highly resolved spatiotemporal platform for studying the cell-cell interactions that can drive resistance to ICIs in diverse TME settings and identifies potential therapeutic avenues to enhance the efficacy of emerging ICI therapies.

## 1 Introduction

Breast cancer is the most prevalent form of cancer in women, and while early-stage breast cancer is associated with high survival rates, metastatic breast cancer continues to have poor prognoses across histologic and molecular subtypes despite advances in targeted therapeutics. Immune checkpoint inhibitors (ICIs) can promote anti-tumor immune responses by blocking inhibitory signaling pathways that inactivate effector T cells. These drugs have seen recent clinical success and are approved in a number of settings, including melanoma and non-small cell lung cancer [1]. In breast cancer specifically, however, the only approved use of ICI is for PD-1 inhibitors in combination with chemotherapy in specific cases of triple-negative breast cancer (TNBC) [2]. In addition, dual checkpoint inhibition, the addition of anti-CTLA-4 to anti-PD-1 therapy, has been efficacious in lung cancer, melanoma, and primary liver cancer; however, its use in breast cancer is still emerging [3]. In a meta-analysis of 10 trials, which included a total of 43/728 patients who received dual ICI, pooled objective response rates (ORR) were 29.4% compared to 7.1% in those who received single-agent ICI (95% CI, 15.7-44.9%). Estimated 6- and 12-month progression-free survival (PFS) rates were 12.2%/5.5% for single ICI and 38.7%/31.4% for dual ICI. Among dual ICI recipients, 21% achieved disease control for >12 months and 14% for >24 months. Notably, a subset of patients can experience durable response [3]. Although the small dual-checkpoint population and indirect cross-trial comparison preclude a definitive conclusion of superiority, the findings support prospective evaluation of dual checkpoint blockade [3]. Therefore, there is strong rationale to better understand the mechanisms driving response to dual ICI, which could inform biomarker evaluation and future clinical trial design.

In addition, observations about checkpoint response by site of disease are reported across numerous single and dual ICI studies; however, little is known about how response and resistance differ across metastatic organs. The clinical relevance of this is that progression at a single metastatic site can determine clinical treatment failure, even when disease elsewhere remains controlled. Under RECIST, new lesions or unequivocal progression of non-target disease can establish overall progression, while aggregate target-lesion measurements can obscure divergent growth and shrinkage across organs [4, 5]. Organ-specific progression therefore provides a clinically consequential readout of treatment escape, while imaging alone cannot distinguish inadequate drug expose, local immune suppression, and tumor-intrinsic resistance. Identifying where and when escape occurs is essential to explaining why an initially active systemic regimen fails. For example, studies emphasize patient-level response and progression-free survival, leaving unresolved whether lung metastases differ in response depth, durability, or timing of immune escape. Biological observations support investigating this question: in a trastuzumab-treated HER2-positive cohort, pulmonary and hepatic metastatic patterns were associated with different primary-tumor immune profiles and outcomes, but these findings did not establish organ-specific checkpoint sensitivity [6]. Mechanistic observations in human melanoma demonstrate that PD-1 and CTLA-4 blockade engage partly distinct T cell populations: CTLA-4 inhibition promotes expansion of ICOS-positive, Th1-like CD4+ effector cells, in addition to effects on CD8+ T cells [7]. Whether these complementary effects improve control of metastases, and whether their contribution differs between lung, liver, and other organs, remains unresolved.

The highly immunosuppressive tumor microenvironment (TME) of breast cancer relative to other solid tumors has been implicated as a contributing factor to the ineffectiveness of ICIs [8, 9]. In this environment, interactions between effector cells and neighboring cells, such as PD-1/PD-L1 interactions, can attenuate effector cell functions, leading to poor control of cancer cell growth even in tumors with high infiltration of CD8+ T or natural killer (NK) cells. Additionally, cancer cells and pro-tumor immune cells such as myeloid-derived suppressor (MDSC)-like cells, T regulatory cells (Tregs), and M2-like macrophages can secrete factors that induce pro-tumor behaviors in immune cell populations that would normally contribute to an anti-tumor immune response. The complexity of the TME complicates detailed *in vivo* studies of its cellular components and their behaviors, and *in vitro* studies are overly simplistic due to their reliance on individual cell types or combinations of a few cell types in isolation. Thus, a detailed understanding of interactions within the TME cannot be accomplished with a purely experimental approach.

Computational models allow researchers to integrate knowledge from *in vitro* and *in vivo* experiments to analyze the TME from a systems perspective. To date, various computational approaches have been applied to the TME, including differential equation-based methods and agent-based methods [10–13]. Agent-based models (ABMs) are especially well-suited for studying the TME, as they can explicitly model the spatial organization of tumors, while accounting for direct and indirect interactions between cells [14–17]. Furthermore, recent ABMs have been initialized directly from multiplexed images of resected tumors, allowing researchers to integrate knowledge about the TME’s spatial organization into their models [18–21]. The spatial organization of a metastasis can be particularly important when considering response in different visceral organs as the constraints vary tremendously depending on location of metastatic tumor growth.

Integrating organ-specific measurements with mathematical modeling offers a way to investigate these processes more efficiently. In metastatic breast cancer, lesion-level models have quantified differences in growth and treatment-induced shrinkage across organs and linked lesion dynamics and new-lesion appearance to survival [5]. Mechanistic modeling can help distinguish how, where, and under what conditions adding anti-CTLA-4 may improve treatment. Fitting these models to longitudinal lesion measurements and paired immune profiling would allow estimation of lung-versus-other-organ response dynamics, simulation of alternative treatment schedules, and identification of the measurements most informative for distinguishing mechanisms. This approach could prioritize experiments and clinical hypotheses more efficiently, while requiring comparative treatment data and experimental validation to establish the specific contribution of CTLA-4 blockade.

Here, we present an ABM to simulate the spatiotemporal dynamics of tumor and immune cells in breast-to-lung metastases in response to anti-PD-1 and anti-CTLA-4 (dual ICIs), building on our prior work on tumor-immune dynamics in cancer [22, 23]. The model predicted the effect of ICIs across a heterogeneous cohort of breast-to-lung metastases and identified drivers of resistance to ICIs via analyses of spatial outputs. These analyses and simulation results then informed alternative therapeutic strategies to enhance the effects of ICIs, and we evaluated the efficacy of the proposed treatment strategies using the ABM. Our results highlight how experimentally-informed computational modeling can be an effective tool for both investigating cancer biology at a systems-level and for discovering and evaluating potential therapies prior to preclinical studies.

## 2 Results

### 2.1 Spatially-resolved model of the TME directly integrates experimental data from a pre-clinical mouse model of metastatic breast cancer

Initial conditions for the model were directly obtained from a syngeneic mouse model of spontaneous breast-to-lung metastases that were used in previous studies to investigate the immunologic mechanisms of response to dual ICIs [24]. Of note, these mice were treated with therapy that parallels one of the studies included in the meta-analysis of clinical trials involving dual ICIs [3, 25]. Briefly, lung metastases were resected and sliced, and then stained with a nine-color antibody panel for multiplexed immunohistochemistry (mIHC) analysis. After obtaining scans of specific regions of interest, an imaging analysis pipeline was used to segment cells, determine their 2D location, calculate the fluorescence values of each color (continuous variables), and decide whether or not a protein was expressed on a given cell (binary variables) (**Fig. 1A**). A subset of cells was labeled using binary protein expression for each of the nine markers based on expected protein expression profiles for eight distinct cell types commonly found in tumors and included in the model: cancer cells, CD8+ T cells, NK cells, CD4+ T cells, Tregs, MDSC-like cells, M1-like macrophages, and M2-like macrophages (**Sup. Table S2**). Any cells with expression profiles that were not associated with the specific cell types included in the model remained unlabeled. Successfully labeled cells were then used to train a random forest model incorporating the continuous fluorescence values obtained by the mIHC pipeline, and this model was used to predict the identities of the unlabeled cells.

**Figure 1:**
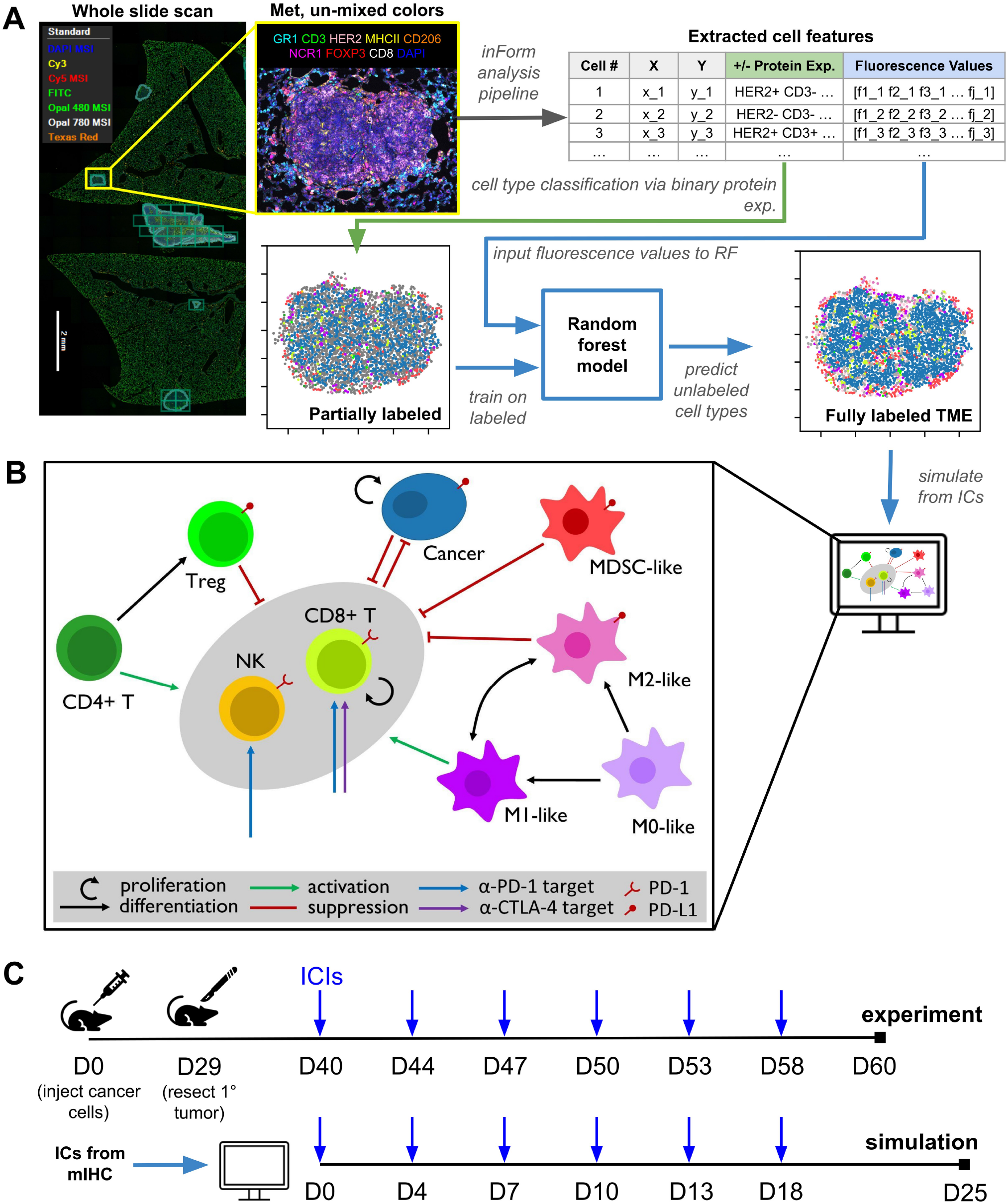
A syngeneic mouse model informs an agent-based model of immune dynamics in the breast-to-lung metastatic tumor microenvironment. **A.** Pipeline for obtaining initial conditions (ICs) from mIHC imaging of mouse breast-to-lung metastases to initialize the agent-based model (ABM). **B.** Summary of cell-cell interactions and immune checkpoint inhibitors present in the ABM. C. Simulated treatment protocol based on an experimental protocol for dual ICIs (anti-PD-1 and anti-CTLA-4) with entinostat administered twice-weekly.

Additionally, the model included three generalized cell types with limited behavior (i.e., they participate in cell physics, and immune lineage cells die over time): myeloid cells, lymphoid cells, and stromal cells. These generalized cell types were assigned to cells that had low-confidence cell type predictions from the random forest model (**Sup. Fig. S1**). While these cells are needed to preserve the overall geometry of the tissue, defining them as generalized cells avoided the assignment of biological behaviors that yield unrealistic model simulations. To prioritize simulation of metastases that are biologically accurate, we excluded metastases with high generalized cell populations. We also excluded metastases with very low cancer cell populations, metastases with high CD8+ T cell and low cancer proportions that could represent tertiary lymphoid structures, and very small metastases (see **Methods**). Following this mIHC pipeline, we selected 19 metastases from an initial population of 80 to simulate with the ABM.

The ABM included several parameters that govern cell dynamics based on literature values, experimental observations, and prior modeling work (see **Methods** and **Supplementary Methods**; **Fig 1B** shows interactions in the model). Additionally, some parameters related to CD8+ T cells were specifically tuned to reflect the TME of breast-to-lung metastases as follows. To identify a parameter value for CD8+ T cell recruitment and to validate calculated parameters in the exhaustion model, we conducted sensitivity analyses and classified the behavior of each metastasis based on the relative number of cancer cells at the study endpoint. These classifications were based on the irRECIST criteria: complete response (CR) indicates complete elimination of cancer cells, partial response (PR) indicates a reduction in cancer cell population by 30% or more, progressive disease (PD) indicates an increase by 20% or more, and stable disease (SD) indicates a change between -30% and +20%, all relative to the initial cancer cell population [26].

In our model, CD8+ T cells are recruited proportionally to the number of cancer cell deaths at a given time step. To determine the rate of CD8+ T cell recruitment, we varied this parameter over a range of values and qualitatively examined the outcomes for four metastases with different initial proportions of cancer cells, pro-tumor immune cells, and anti-tumor immune cells (**Sup. Fig. S2A**). Two of these metastases, Mets 30 and 7, had low initial CD8+ T cell populations (0.6% and 7.1%, respectively). The other two metastases (Mets 0 and 34) had higher initial CD8+ T cell populations (19.9% and 56.1%, respectively). Since breast cancer metastases generally lead to death in the experimental mouse model, we selected a CD8+ T cell recruitment rate for which a substantial number of replicates were classified as PD, rather than SD.

We also characterized the effect of parameters related to the rate of CD8+ T cell exhaustion. To do this, we varied the exhaustion property-specific multiplicative factor, *k_d_*, to study how the rate of exhaustion for the probability of killing cancer cells (P(kill)), the cell cycle length, and the cell lifespan influenced the outcome of the lung metastases simulated by the ABM (**Sup. Fig. S2B, C**). For higher values of the property-specific *k_d_* (corresponding to slower exhaustion), we observed decreases in the final cancer population for all metastases when considering the *k_d_* values associated with P(kill) and the cell cycle length. These trends were preserved when simulating with anti-PD-1 and anti-CTLA-4 dual ICI treatment, with slight reductions in the final cancer cell population across all parameter values and metastases. Varying the *k_d_* value for the CD8+ T cell lifespan did not meaningfully change the distribution of final cancer cell populations, even when setting the parameter value to 1, which prevents cell interactions from lowering the cell’s lifespan. With treatment, certain metastases had overall reductions in cancer growth, but the trend remains the same.

Finally, we varied the baseline probability of a CD8+ T cell killing a cancer cell, P(kill), after making direct contact. For the metastases with higher initial CD8+ T cell populations, increasing this value led to consistent decreases in the final cancer cell population. In fact, increasing the probability to 0.4 led to CR in all replicates in the metastasis with high initial CD8+ T cell count (Met 34). However, there was no meaningful decrease in final cancer cell population in the two metastases with low initial CD8+ T cell populations after increasing the probability from 0.2 to 0.4. Conducting this analysis while simulating dual ICIs led to the same trends, but with slightly lower values for final cancer cell populations.

Overall, we used these sensitivity analyses to explore the behavior produced by the model and to select appropriate model parameters. We specified the CD8+ T cell recruitment rate and confirmed that the values related to CD8+ T cell probability of killing, cell cycle length, and lifespan produce simulations where the outcome based on cancer cells aligned with preclinical observations.

### 2.2 The simulated dynamics of cell counts vary widely across cell types and metastases

After initializing the ABM with mIHC data following the steps outlined in **Fig. 1A**, we simulated the dynamics of each of the 19 metastases for 600 hours without and with dual ICIs, conducting 25 replicates per metastasis with the treatment schedule in alignment with pre-clinical mouse studies (**Fig. 1C**). The simulation results for the no treatment case are shown in **Fig. 2**. For the majority of metastases (all except Mets 13, 32, and 42), we observed two distinct regimes of dynamics: an initial period (to 200 hours) where there were relatively large and rapid changes in the cell populations, and a subsequent period where some metastases appeared relatively stable while other metastases grew at slow, linear rates. The metastases that did not follow this trend still often featured very short-term initial changes in their macrophage, CD4+ T cell, and Treg populations, but had a longer, less noticeable transition between the two dynamical regimes after these initial changes.

**Figure 2:**
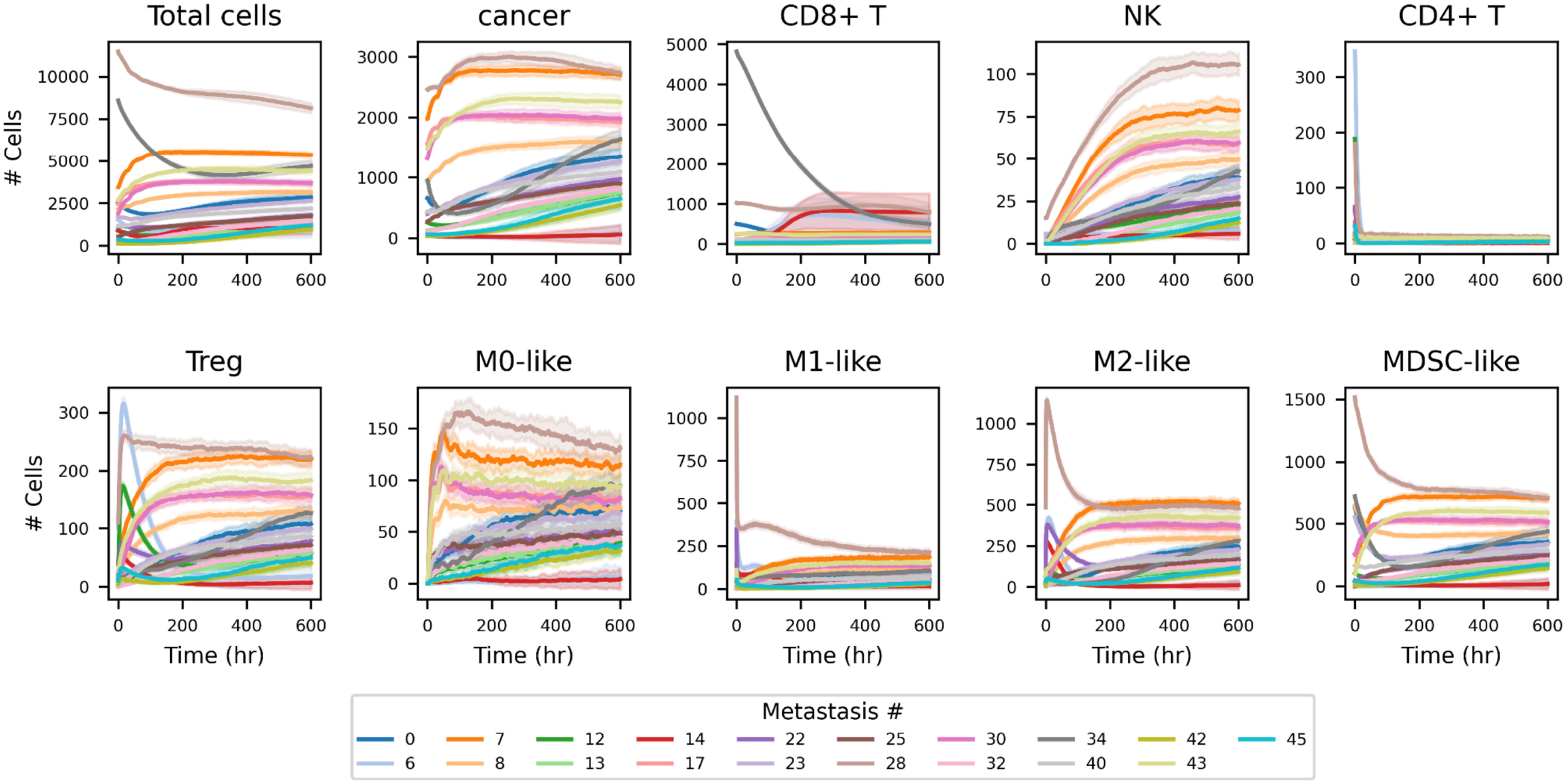
Simulated bulk cell count dynamics for 19 metastases without treatment reveal heterogeneous behaviors. Dynamics are shown for all nine functional cell types and the total number of cells. The total cell count panel (top left) includes the three generalized cell types (myeloid, lymphoid, and stromal).

Metastases with larger initial cancer cell populations tended to have more drastic changes in the first 200 hours and were more likely to exhibit stable dynamics relative to those that began with lower cancer cell populations. However, this was not always the case. For example, Mets 8 and 34 started with similar cancer cell populations but only the dynamics of Met 8 stabilized, while Met 34 continuously expanded following an initial period of decline. Additionally, we observed that the cell types with the most dramatic initial dynamics were involved in differentiation or re-polarization processes influenced by the local microenvironment, including CD4+ T cells, Tregs, and M1- and M2-like macrophages. In general, the initial dynamics for these cell populations were dominated by the conversion of anti-tumor cell types (CD4+ T cells, M1-like macrophages) into pro-tumor cell types (Tregs, M2-like macrophages).

### 2.3 Modeling predicts substantial heterogeneity in the endpoint cancer cell populations across metastases

After simulating the dynamics of all cell types in each metastasis, we classified the endpoint cancer cell population based on the irRECIST criteria (populations normalized to initial conditions). At the end of the simulation window, most metastases displayed PD with or without treatment for all replicates (73%), while a small subset of metastases (16%; Mets 34, 17, and 28) had some replicates classified as SD with treatment (**Fig. 3**). Of these metastases, only Mets 17 and 28 had replicates that were classified as SD even in the absence of treatment. Further, only two metastases (11%; Mets 6 and 14) had replicates that reached CR. However, simulations for these metastases also had replicates that were classified as PD, SD, or PR. Thus, these metastases displayed significant heterogeneity across replicates, compared to other metastases whose endpoint cancer cell populations were confined to relatively smaller ranges.

**Figure 3:**
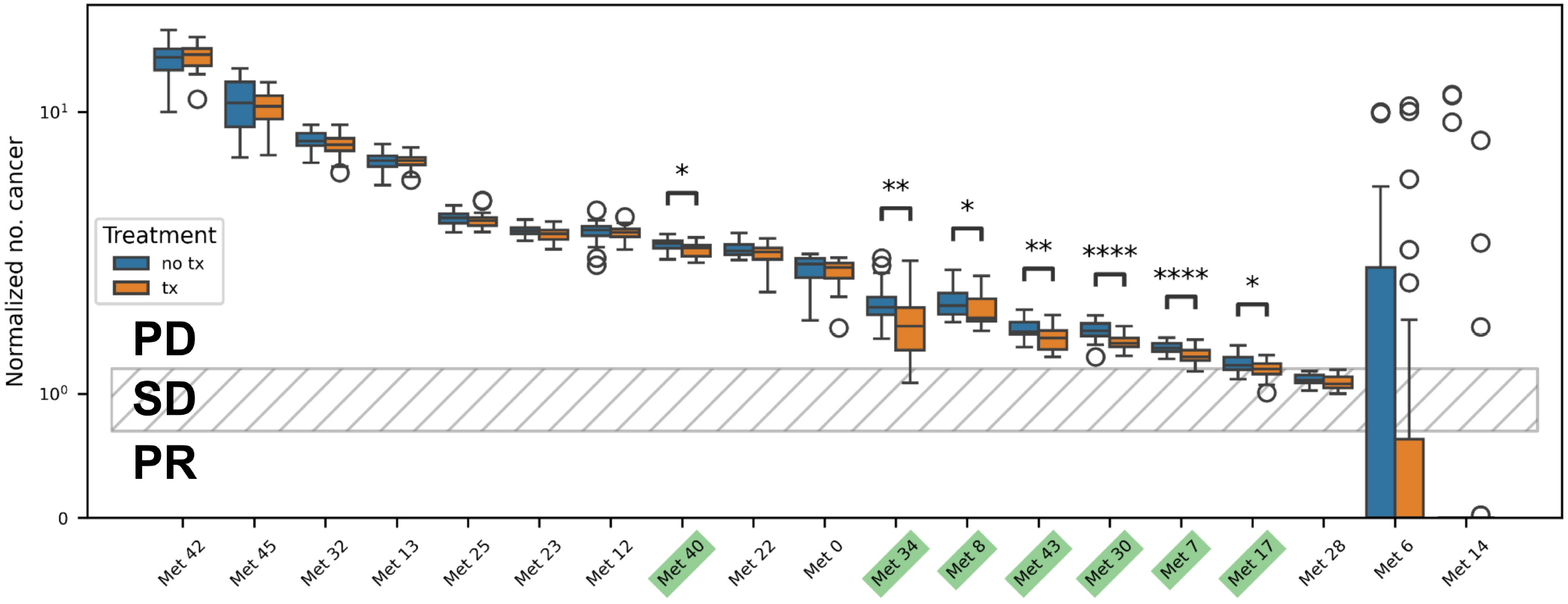
Classification of metastases into irRECIST-based categories identifies metastases for which dual ICIs have an effect. Final cancer cell count is normalized to the initial cancer cell count. Green labels and asterisks indicate metastases for which there is a statistically significant difference between treatment and no-treatment groups, as evaluated using either a student’s *t*-test of a Mann-Whitney *U* test: * p *≤* 0.05, ** p *≤* 0.01, *** p *≤* 0.001, **** p *≤* 0.0001.

Given this significant intra-metastasis heterogeneity, we separately studied Mets 6 and 14. These metastases had a high percentage of immune cells: 80% and 67%, respectively (**Sup. Fig. S3A**), compared to an average of 38% for the other metastases shown in **Fig. 2**. We performed an additional 75 simulations for these two metastases to study the heterogeneous outcomes and found that most replicates were classified as CR without treatment (46% and 93% for Met 6 and Met 14, respectively **Sup. Fig. S3B**). Across both metastases, replicates not classified as CR exhibited a wide range of growth patterns, potentially influenced by the higher proportions of immune cells and their cellular interactions.

In Met 6, 36% of the replicates were classified as PD, which could be be grouped into two categories based on their cancer cell expansion trajectories: one group grew more rapidly and became distinguishable from the CR replicates before 300 hours, while the other group only became distinguishable after 400 hours and achieved lower growth. For Met 14, the seven replicates that did not achieve CR are classified as PD (6) or PR (1). Simulation results showed that these replicates either entered a linear-like growth phase prior to 200 hours or grew at a slower rate and were indistinguishable from the main cohort of CR replicates until after 200 hours. For both metastases, applying dual ICIs led to visible reductions in cancer cell expansion for the replicates classified as PD. In Met 6, treatment additionally led to fewer simulations classified as PD. In contrast, for Met 14, one replicate classified as CR was instead classified as PR, while the number of replicates that achieved PD remained the same. Altogether, we concluded that these metastases displayed divergent outcomes that could be influenced by dual ICIs.

For the 19 metastases we simulated, only seven (37%; Mets 7, 8, 17, 30, 34, 40, and 43) demonstrated statistically significant decreases in the mean endpoint cancer cell population with treatment. Dual ICIs did not affect cancer cell population growth in the remaining 12 metastases. Relative to the complete cohort, the metastases in which dual ICI treatment had an effect also had lower normalized endpoint cancer cell populations (excluding Met 40). Additionally, the decline in mean cancer cell growth for metastases affected by simulated treatment ranged from 3.7% to 10.4%, and only Met 17 had its average outcome move from PD to SD with the addition of dual ICIs.

Despite dual ICIs not having a clinically relevant effect on cancer cell growth, model simulations confirmed that dual ICIs altered the behavior of CD8+ T cells within the TME. Since we modeled CD8+ T cell exhaustion as a continuous process, we were able to extract and visualize the trajectories of exhaustion-related parameters for each CD8+ T cell within a single replicate to understand how the TME influenced CD8+ T cells’ ability to control growth of the cancer cell population. As an example, **Sup. Fig. S4A** shows the probability of CD8+ T cells killing a cancer cell upon making direct contact for each CD8+ T cell in Met 0 over time. This analysis showed high heterogeneity in the killing probability feature of CD8+ T cell exhaustion. Interestingly, some CD8+ T cells retained a relatively high killing probability even 200 hours after their birth. We integrated these trajectories to produce a single “score” that reflects the total potential efficacy of a CD8+ T cell over its lifetime. Considering the score for each CD8+ T cell across replicates in a metastasis yielded a distribution of scores that provided insight into how these cells function in the metastasis. The distribution of this metastasis-level score for CD8+ T cell killing is shown in **Sup. Fig. S4B** for all metastases except Met 6 and Met 14 (due to their divergent outcomes, as described above). We found that there was a large exponentially-distributed population of CD8+ T cells with low integrated scores, which may include cells representing the progeny of already-exhausted CD8+ T cells, cells that rapidly become exhausted, or cells that quickly die off. Additionally, there was another large population of CD8+ T cells with normally-distributed integrated scores. The means of these two distinct CD8+ T cell populations tended to be lower in metastases for which dual ICIs had an effect on outcome. Thus, these scores allowed us to directly observe the effect of dual ICIs on killing probability in CD8+ T cells (**Sup. Fig. S4C**), as the medians of the normally-distributed cell population were significantly elevated in the treatment groups for all metastases except for Mets 34 and 42. These elevated scores indicate that the checkpoint inhibitors slowed the rate of exhaustion.

### 2.4 Metastases for which dual ICIs have an effect display similar patterns of cell-cell co-localization

We next aimed to understand whether spatial features contribute to the effect of dual ICIs. To do so, we first studied co-localization patterns of pairs of cells. Calculating the co-localization Z-score for each pairwise combination of cell types allows us to summarize the spatial organization of each metastasis in a quantitative fashion (i.e., as a matrix), which can then be projected into 2D space via a dimensionality reduction technique such as UMAP (**Fig. 4A**; the UMAP was calculated on the 1D vectors formed from the upper triangular entries of each Z-score matrix). When we compared the co-localization Z-score matrices for metastases over time, we saw that the co-localization of pairs of cell types changed over time. That is, the spatial organization on a cell-cell pairwise basis was not static (**Fig. 4B**). We observed that positions of the initial states (yellow color) of metastases for which dual ICIs had an effect sometimes overlapped with metastases for which treatment did not have an effect. However, the final states of metastases (dark purple color) with and without a treatment effect are mapped to different regions of the manifold. These mappings illustrate that while metastases with and without treatment effect may begin with similar cell-cell co-localization scores, the metastases for which treatment had an effect move to distinct regions of the UMAP, distinguishing these metastases from the others. In contrast, the metastases for which treatment does not have an effect displayed less noticeable changes in their location in the UMAP over time. More broadly, we can conclude that the spatial organization of a metastasis is correlated with whether or not dual ICIs have an effect.

**Figure 4:**
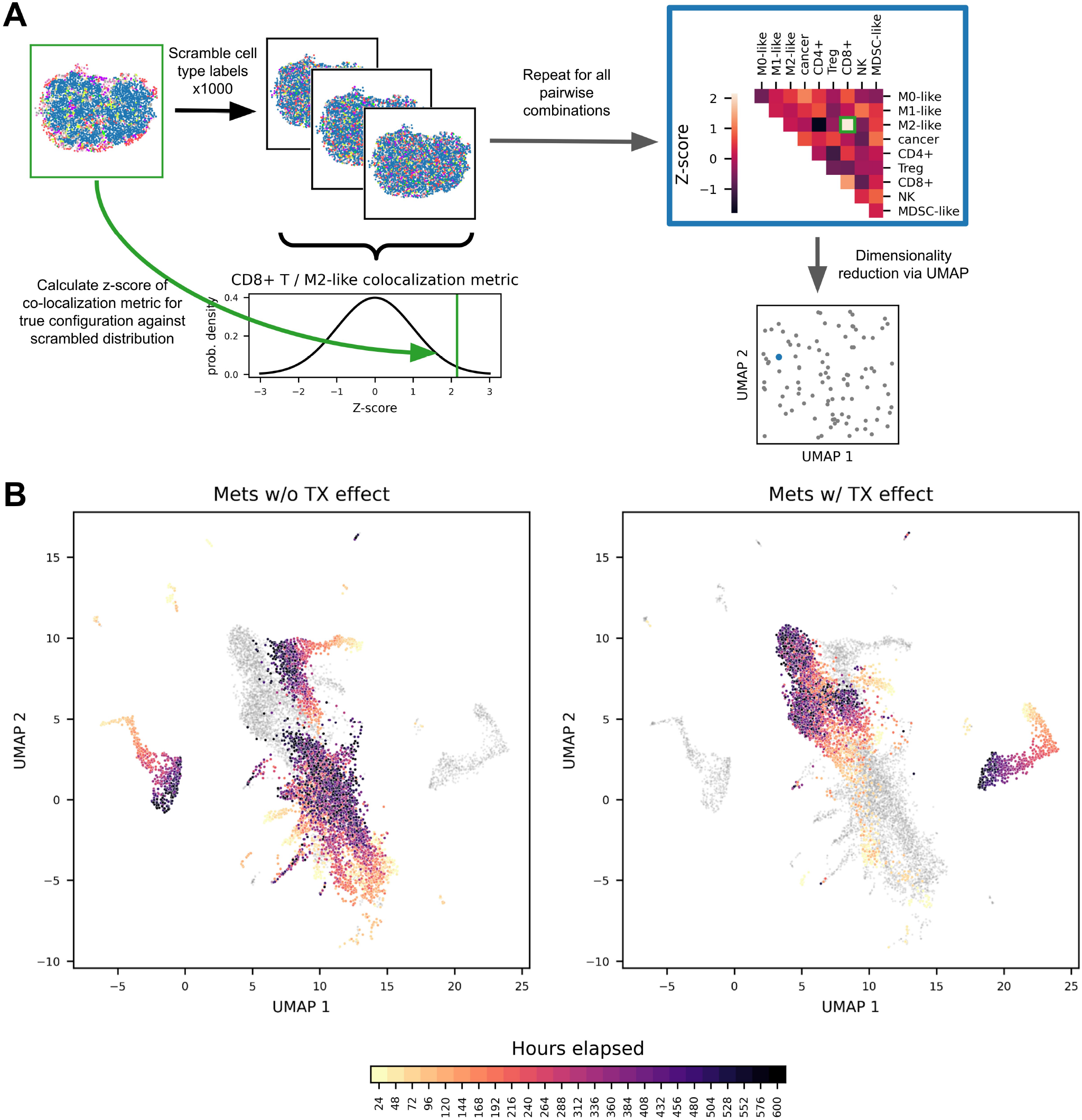
Two-dimensional UMAP of the neighborhood enrichment Z-score matrices reveal structural similarities for metastases in which dual ICI treatment has an effect. **A.** Pipeline for producing 2D representations of neighborhood enrichment Z-score matrices. **B.** UMAP of the spatial configurations of metastases sampled every 24 hours, excluding replicates that reach CR before the sampled time. Metastases were separated based on whether dual ICI treatment causes a statistically significant reduction in cancer growth (Left: Mets 0, 12, 13, 22, 23, 25, 28, 32, 42, and 45; right: Mets 7, 8, 17, 30, 34, 40, and 43).

### 2.5 The presence of cell triplets involving CD8+ T cells reflects the anti- and pro-exhaustion regions in simulated metastases

As metastases are composed of more than two cell types, we moved beyond studying pairwise co-localization of cells to investigating the presence of triplets of cells. We particularly considered triplets that include CD8+ T cells to understand how specific immune cell types affect the CD8+ T cell exhaustion phenotype. For simulated spatial outputs, we constructed a pruned Delaunay triangulation to identify potential cell interactions, which were filtered to retain cell interactions that were either direct contacts or strong indirect interactions. From these remaining edges, we quantified the number of CD8+ T cells with cancer cell neighbors, as these cells are the population of CD8+ T cells that could potentially kill cancer cells at any given time point. We then quantified the effect of other immune cells (including other CD8+ T cells) by counting the number of CD8+ T cells with cancer cell neighbors that also had an immune cell neighbor. An example of such a triplet is shown in **Fig. 5A** for a CD8+ T cell having both a cancer cell and MDSC-like cell as neighbors. We call the ratio of such triplets to cancer cell-CD8+ T cell dyads the “interaction ratio” (IR). The IR is a meaningful quantity to consider, as the interaction of CD8+ T cells with cancer cells directly regulates the growth of the cancer cell population, and the interactions of CD8+ T cells with other immune cells have a direct impact on the cell-intrinsic parameters that control a specific CD8+ T cell’s behavior in simulation through our exhaustion model. We then calculated the IRs for all immune cell types (for the no-treatment simulations).

**Figure 5:**
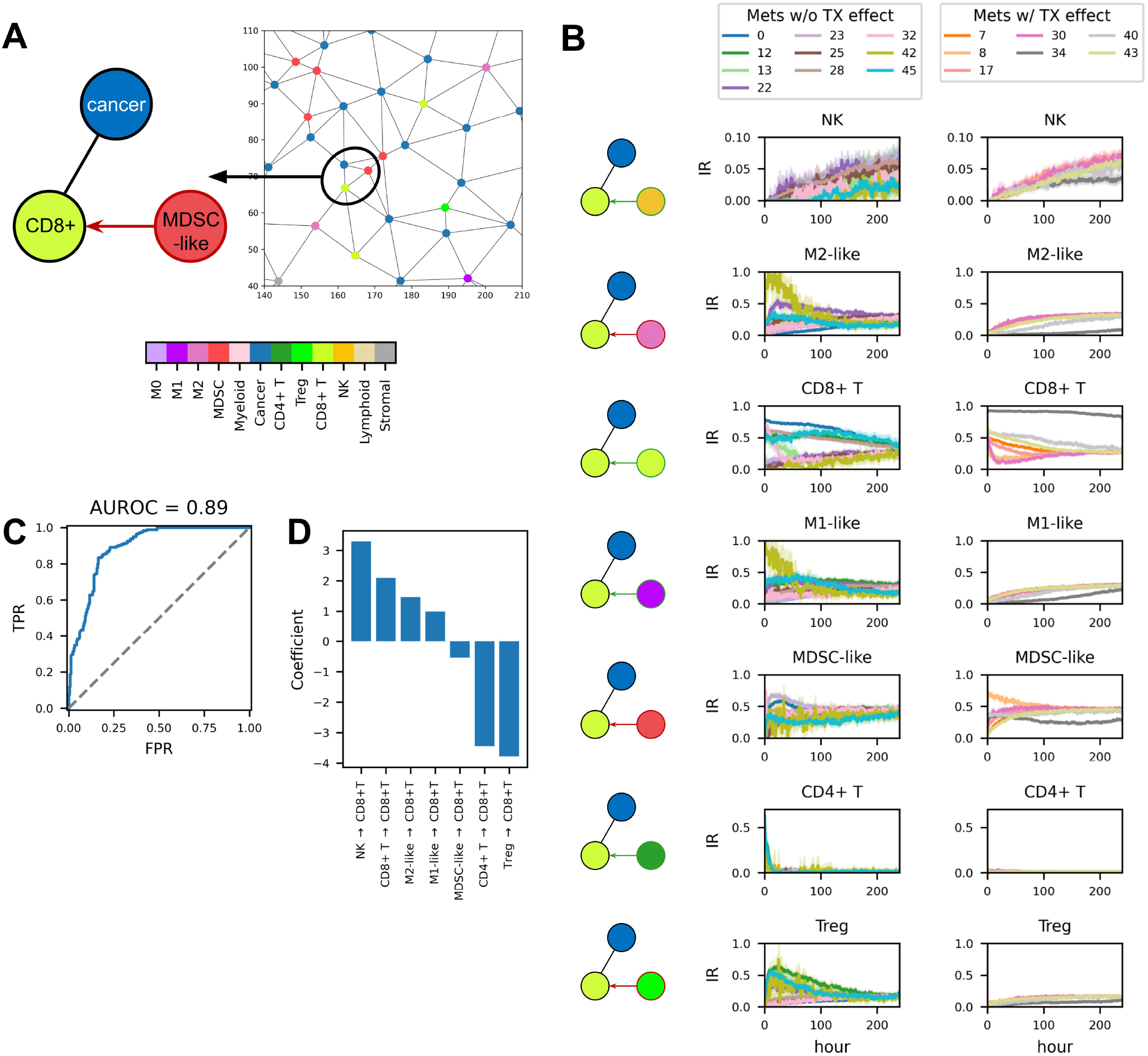
Spatially-constrained cell-cell interaction scores can predict treatment effect via logistic regression. **A.** Overview for extracting cell triplets from the pruned Delaunay triangulation of an ABM spatial output. **B.** Spatially-constrained cell-cell interaction ratios (IRs) over time for metastases without treatment effect (left column) and with treatment effect (right column). The IR (y-axis) is defined as the number of triplets for a given cell type over the total number of CD8+ T cell-cancer cell dyads, and thus ranges from 0 to 1. Note that the axes for the NK cell and CD4+ T cell IR are not from 0 to 1 in order to highlight differences between groups. **C.** Receiver operating characteristic (ROC) curve for a logistic regression model with time-integrated IRs as inputs and prediction of whether treatment will have an effect as the output. **D.** Logistic regression coefficients for the same model. Positive coefficient - input is associated with treatment effect; negative coefficient - the input is associated with treatment resistance.

We observed that for most immune cells, the IR approached a constant value by 240 hours in most metastases with no treatment effect. However, the initial dynamics of the IR scores varied across metastases (**Fig. 5B**). Values for the NK and CD4+ T cell IRs were low compared to other immune cells, indicating that interactions between CD8+ T cell-cancer cell dyads and NK or CD4+ T cells were less prevalent than for other immune cells. We also observed that metastases with a treatment effect generally had similar IR trajectories (except Met 34). In comparison, metastases without a treatment effect demonstrated more variability across IRs for each cell type.

We next aimed to understand whether these interactions between triplets of cells inform whether a metastasis is influenced by dual ICIs. We fit a logistic regression model to predict whether dual ICI treatment would affect the relative growth of the cancer cell population.

We used the integrated IR (area under the IR curve) for each immune cell as inputs to the regression model, termed “Regression Model 1.” Thus, this regression model has seven total inputs, which were each min-max normalized across simulations. After determining the optimal number of days to use in calculating the integrated IR (see **Methods**), we fit the logistic regression to the complete data set containing all no-treatment simulation replicates from the ABM. The AUROC score for this model was 0.89 (**Fig. 5C**), and both the accuracy score and the balanced accuracy score were 81.1%. These metrics indicate that the presence of specific triplets can capture whether dual ICI treatment will have an effect on the outcome for individual metastases.

By analyzing the regression coefficient associated with each integrated IR score, we can determine which cell-cell interactions most strongly influence whether dual ICI treatment is effective in a given metastasis (**Fig. 5D**). A positive coefficient means the input is associated with treatment effect, while a negative value indicates the input is associated with treatment resistance. For this regression model, the NK cell IR was most strongly associated with treatment effect, with a regression coefficient of 3.3. This is followed by the CD8+ T cell IR, the M2-like macrophage IR, and the M1-like macrophage IR (regression coefficients of 2.1, 1.5, and 1.0, respectively). The input most associated with treatment resistance was the Treg IR, followed by the CD4+ T cell IR, and the MDSC-like cell IR (regression coefficients of -3.8, -3.4, and -0.5, respectively). While CD4+ T cells are considered anti-tumor cells in our model, as they can enhance CD8+ T cell functionality, model simulations showed that CD4+ T cells typically differentiated into Tregs in early stages of simulation. This provides a possible explanation for the counterintuitive finding that integrated IRs for CD4+ T cells predict resistance to dual ICIs.

We also trained logistic regression models that included multiplicative, pairwise combinations of integrated IRs to evaluate whether there were interactions between IRs of two cell types that would enhance the regression model’s ability to discriminate between metastases with treatment effect and those without. First, we evaluated the performance of a regression with the original inputs in addition to specific pairwise combinations (“Regression Model 2”), totaling 24 inputs. These combinations included all pairwise combinations of pro-tumor cells with other pro-tumor cells, anti-tumor cells with other anti-tumor cells, and all cells with the cancer vulnerability score, which is the ratio of cancer cells with CD8+ T cell neighbors to total cancer cells (**Sup. Fig. S5A**). For Regression Model 2, the Treg and CD4+ T cell IRs were again the inputs most strongly associated with treatment resistance (coefficients of -2.2 and -1.9, respectively). However, the feature most strongly associated with treatment effect was the pairwise combination of the M2-like macrophage and MDSC-like cell IRs, followed by the NK cell IR (coefficients of 2.0 and 1.8, respectively). The accuracy and balanced accuracy scores of Regression Model 2 were 83.5%. This represents a slight improvement relative to Regression Model 1 (+2.4%) with the addition of 17 input features.

Lastly, Regression Model 3 included all pairwise combinations of IRs, combinations of IRs with cancer vulnerability, and the IRs alone, for a total of 36 inputs. The feature most correlated with treatment effect was the CD8+ T cell IR, with a regression coefficient of 2.4. The feature most correlated with lack of treatment effect was the combination of the MDSC-like IR and cancer vulnerability score, with a regression coefficient of -3.2 (**Sup. Fig. S5B**). Interestingly, the association with treatment was reversed for a number of inputs, compared to Regression Models 1 and 2. After including 36 total features, the regression model’s accuracy score was 84.5% and the balanced accuracy score was 86.0% (improvements of 3.4% and 4.9%, respectively, compared to Regression Model 1).

Analyzing the receiver operating characteristic curves for all three regression models illustrates their relative performance (**Sup. Fig. S5C**). For Regression Models 2 and 3, the AUROC score was only marginally better than Regression Model 1 (both were 0.90, versus 0.89 for Regression Model 1). This result implies that despite the addition of 17 and 29 input features, respectively, these models did not have better performance than Regression Model 1. Thus, the outcomes for ICI treatment effect simulated by the ABM can be well-explained by examining just the integrated IRs. We conclude that NK cells interacting with other CD8+ T cells most strongly correlated with dual ICI treatment having an effect. In contrast, CD8+ T cells interacting with Tregs strongly correlated with a lack of treatment effect.

### 2.6 The relative abundance of specific cellular neighborhoods distinguishes metastases for which ICI treatment has an effect

In addition to studying the interactions between pairs and triplets of different cell types, we also considered the larger neighborhoods that cells occupy. These neighborhood labels, which are calculated based on the cell types present near a given cell, provide an alternative way to describe the spatial organization of a metastasis (**Fig. 6A**). As the ABM simulates the spatial organization of metastases over time, we were able to calculate the size of each neighborhood over time. For example, in the single no-treatment replicate of Met 7 displayed in **Fig. 6B**, the sizes of the MDSC-like enriched tumor neighborhood (neighborhood C) and MDSC-like/Treg-enriched tumor neighborhood (neighborhood G) expand over time, while the cancer cell neighborhood that is not infiltrated by immune cells (neighborhood A) shrank. We calculated neighborhood labels using all cells present in the first 48 hours of the 25 no-treatment replicates of all metastases. We then used these neighborhood labels to investigate how the initial spatial organization of a metastasis might influence whether dual ICI treatment has an effect. To do this, we calculated the fraction of cells assigned to each neighborhood on a per-metastasis basis and performed hierarchical clustering based on the initial composition (**Fig. 6C**). Unsupervised hierarchical clustering demonstrates that metastases in which dual ICI treatment had an effect had similar initial neighborhood compositions. As shown by the green box, we identified a single dendrogram branch that contains each of these metastases except for Met 34, which is an outlier dominated by a large CD8+ T cell region and remains on an isolated branch until the final merge. This branch also contained two metastases for which dual ICI treatment did not have an effect, Mets 25 and 32. The heatmap demonstrates that the uninfiltrated tumor neighborhood (neighborhood A), which is composed of cancer cells with few immune neighbors, dominated in metastases in this branch. On the other hand, most metastases for which treatment did not have an effect contained fewer cells in this neighborhood. This heatmap further demonstrates that metastases for which treatment did not have an effect have higher fractions of myeloid, myeloid-enriched, Treg-enriched, and generally suppressive tumor neighborhoods (neighborhoods D, E, F, G, H, and J), which were less prevalent in metastases for which dual ICIs had an effect.

**Figure 6:**
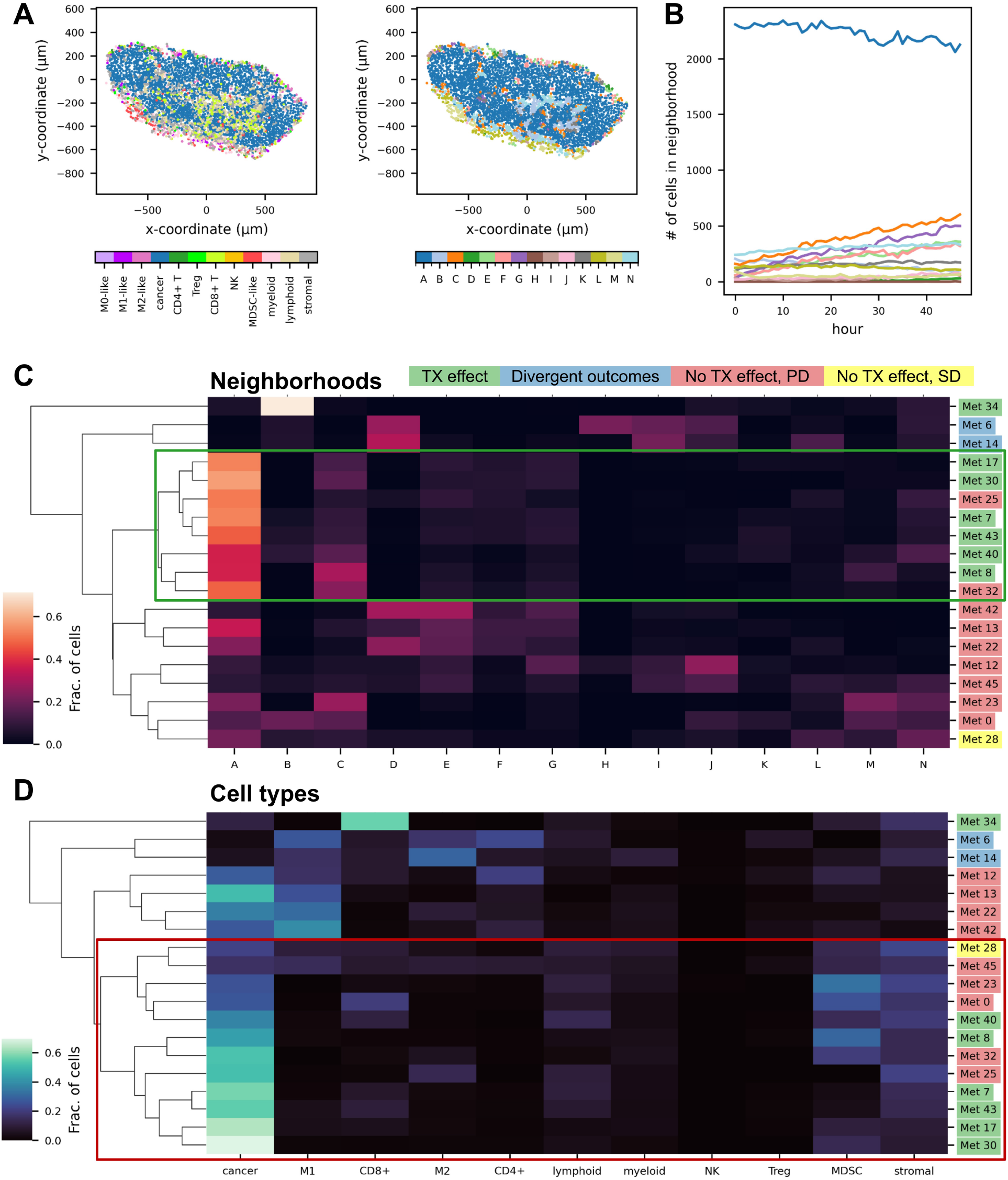
Multicellular structures identified via local neighborhood analysis distinguish metastases by treatment effect. **A.** The spatial organization of Met 7 at time zero with cells colored by cell type (left) and cellular neighborhood (right). **B.** First 48 hours of cellular neighborhood composition for Met 7. The full names associated with each alphabetical neighborhood label are given below. **C.** Heatmap of initial metastasis composition based on cellular neighborhoods. Rows are connected by a dendrogram showing their unsupervised hierarchical clustering. The green box indicates the dendrogram branch containing all metastases for which dual ICIs have an effect except for Met 34. **D.** Heatmap of initial metastasis composition based on cell types. Rows are connected by a dendrogram showing their unsupervised hierarchical clustering. The red box indicates the dendrogram branch containing all metastases that have a treatment effect except for Met 34. **Alphabetical neighborhood labels:** A - uninfiltrated cancer cell region; B - CD8+ T cell region; C - MDSC-like-enriched tumor; D - M2-like macrophage region; E - M2-like macrophage/MDSC-like-enriched tumor; F - M2-like macrophage-enriched tumor; G - MDSC-like/Treg-enriched tumor; H - Treg-enriched tumor; I - T cell activation region; J - Suppressed CD8+ T infiltrated tumor; K - CD8+ infiltrated tumor; L - myeloid region; M - MDSC region; N - generalized cell zone.

We also performed hierarchical clustering of the metastases based only on their initial cell type composition to determine whether spatial neighborhoods explained differences in treatment effect better than bulk cell type proportions (**Fig. 6D**). Using cell-type-based clustering, the delineation between metastases in which treatment did or did not have an effect was less clear than clustering based on cellular neighborhoods. For example, to identify a single branch containing all metastases for which treatment has an effect excluding Met 34 (which clustered on its own as in the neighborhood clustering), we would have to include six metastases for which treatment did not have an effect (red box in **Fig. 6D**). We demonstrate this in greater detail in **Sup. Fig. S6**, which reproduces the dendrograms shown in **Fig. 6C, D** with the maximum distance of each dendrogram normalized to 1. For neighborhood-based clustering, the branch containing all metastases with a treatment effect excluding Met 34 merged at a normalized distance of 0.28 with a branch purity of 75% (the proportion of the branch that was metastases that had a treatment effect). However, the corresponding branch identified via cell-type-based clustering merged at a normalized distance of 0.60 with a purity of 50%. This result shows that the initial spatial organization provides more information about whether or not a metastasis will respond to treatment compared to bulk cell count alone. Additionally, clustering via neighborhoods more effectively isolated metastases without treatment effect, despite their heterogeneity.

### 2.7 Blocking specific cell populations from the TME alters cancer growth and enhances the effect of dual ICIs in certain metastases

Our spatial analyses suggested that the presence of T regulatory cells and MDSC-like cells in certain neighborhoods in the TME contributed to resistance to simulated dual ICIs. Therefore, we evaluated whether blocking these populations from the TME would have a significant impact on cancer growth with and without dual ICI treatment. To model the blockade of MDSC-like cells, we prevented their recruitment into the TME. To model the blockade of Tregs, we prevented CD4+ T cells from differentiating into Tregs. For both cases, we also changed the cell type being blocked into its corresponding generalized immune cell in the initial spatial configurations of the simulated metastases. That is, MDSC-like cells were converted to myeloid cells, and Tregs were converted to lymphoid cells at the start of the ABM simulation.

When blocking MDSC-like cells, only Met 34 had a statistically significant decrease in cancer growth relative to no treatment (growth decreased by 70%; **Fig. 7A**). In fact, most metastases (74%) exhibited statistically significant increases in growth. This is shown in **Fig. 7B**, which summarizes the effect of blockade of MDSC-like cells alone or combined with dual ICIs, compared to no treatment. When combined with dual ICIs, the blockade of MDSC-like cells led to a decrease in the number of cancer cells relative to no treatment for Mets 0, 28, and 34. For Mets 28 and 34, this combination was also an improvement over ICI alone. Additionally, all replicates of Met 14 were CR. However, the combination of ICI and MDSC-like cell blockade led to increased growth in non-CR replicates of Met 6, and did not increase the number of CR replicates relative to ICI alone.

**Figure 7:**
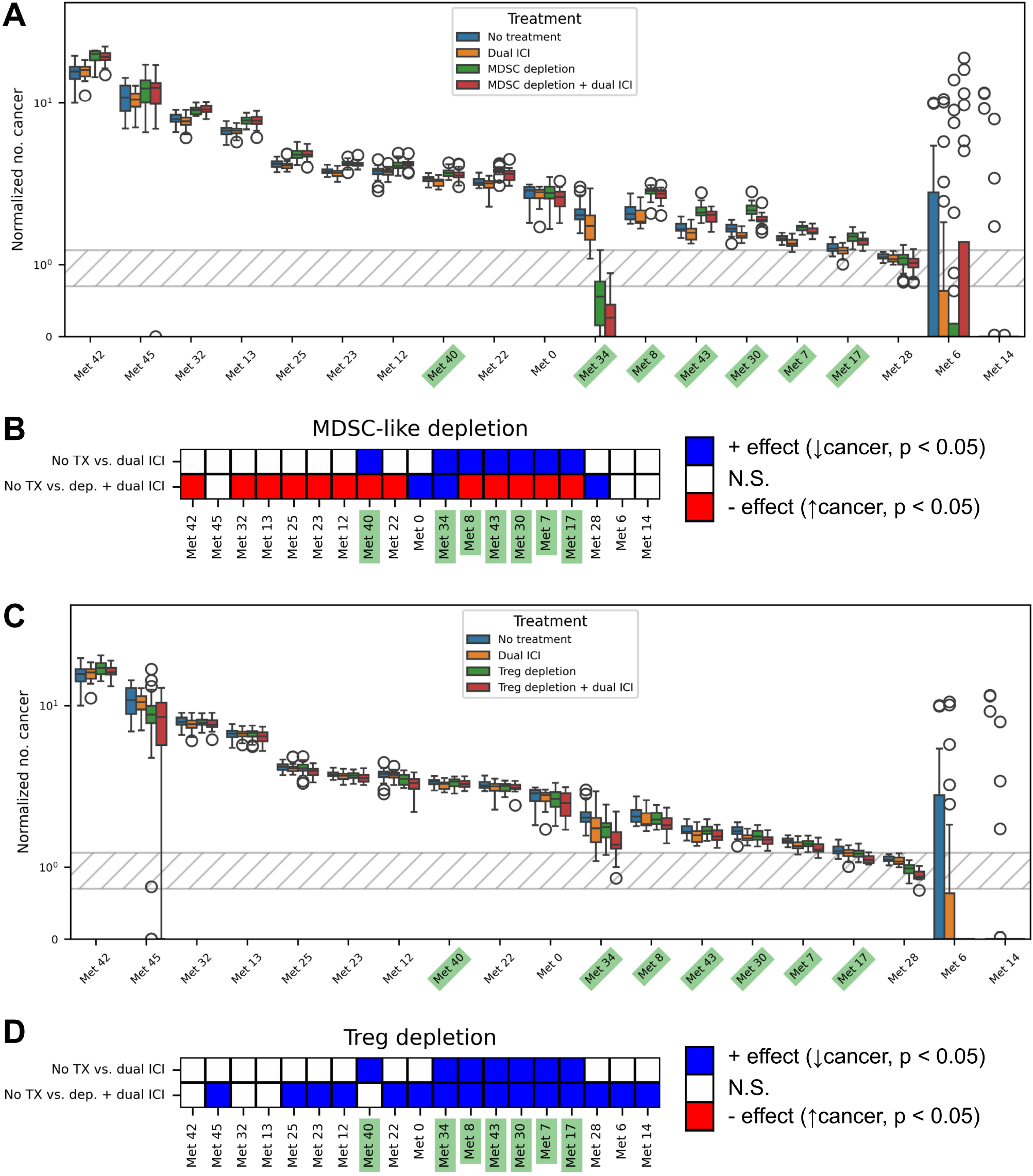
Blocking select cell populations from the TME influences cancer cell proliferation and the effectiveness of immune checkpoint inhibition. **A.** Boxplot showing the distribution of endpoint cancer cell populations (normalized number of cancer cells) for four groups: no treatment, dual ICIs alone, MDSC-like cell blockade alone, and MDSC-like cell blockade with dual ICIs. **B.** Heatmap illustrating the efficacy of dual ICIs alone (top row) or dual ICIs plus MDSC-like cell blockade (bottom row) across metastases compared to no treatment. Red indicates a statistically significant increase in the final cancer cell population, while blue indicates a statistically significant decrease. **C.** Boxplot showing the distribution of endpoint cancer cell populations for four groups: no treatment, dual ICIs alone, Treg blockade alone, and Treg blockade with dual ICIs. **D.** Heatmap illustrating the efficacy of dual ICIs alone (top row) or dual ICIs plus Treg blockade (bottom row) across metastases compared to no treatment. Red indicates a statistically significant increase in the final cancer cell population, while blue indicates a statistically significant decrease. For all panels, green labels indicate metastases for which there is a statistically significant difference between no treatment and dual ICIs alone.

Blocking Tregs alone led to statistically significant decreases in cancer growth for nine out of the 19 metastases (47%), compared to no treatment (**Fig. 7C**). Combining Treg blockade with dual ICIs led to statistically significant decreases in cancer growth for 15 of the 19 metastases (80%), as shown in **Fig. 7D**. For nine of these metastases, the response to combining Treg blockade and dual ICIs was statistically significantly better compared to ICI alone. In fact, all replicates of Mets 6 and 14 reached CR. Thus, Treg blockade decreases the number of cancer cells and can enhance the effect of dual ICIs.

Lastly, we explored how differences in the distribution of MDSC-like cells compared to CD8+ T cells explain why targeted blockade of Tregs enhances dual ICIs. We consider Met 12 as an example, a metastasis that is not affected by dual ICI treatment alone, compared to no treatment (**Fig. 7B**). For Met 12, combining dual ICIs with the blockade of MDSC-like cells leads to growth of the cancer cell population compared to no treatment. In contrast, combining dual ICIs with Treg blockade reduces the size of the cancer cell population compared to no treatment (**Fig. 7D**). The initial spatial organization of this metastasis (**Fig. 8**, middle panel) shows both MDSC-like cells and Tregs. From this, one may expect that blockade of either cell type would lead to reduced growth of the cancer cell population. However, the majority of the MDSC-like cells are positioned away from anti-tumor immune cells. Furthermore, spatial analysis indicates the presence of neighborhood J (**Fig. 8**, right panel), comprised of CD8+ T cells co-localized with Tregs and CD4+ T cells, which rapidly differentiate into Tregs in pro-tumor environments. These differences in initial spatial organization help explain the differential effects of targeted blockade of a particular immune cell type. Specifically, as Tregs are more closely associated with CD8+ T cells, blocking Tregs reduces immunosuppressive effects on the CD8+ T cells, permitting cancer cell killing. This analysis further emphasizes that the effect of immunotherapy depends on the spatial neighborhoods in the metastasis, rather than simply whether a particular cell type is present.

**Figure 8:**
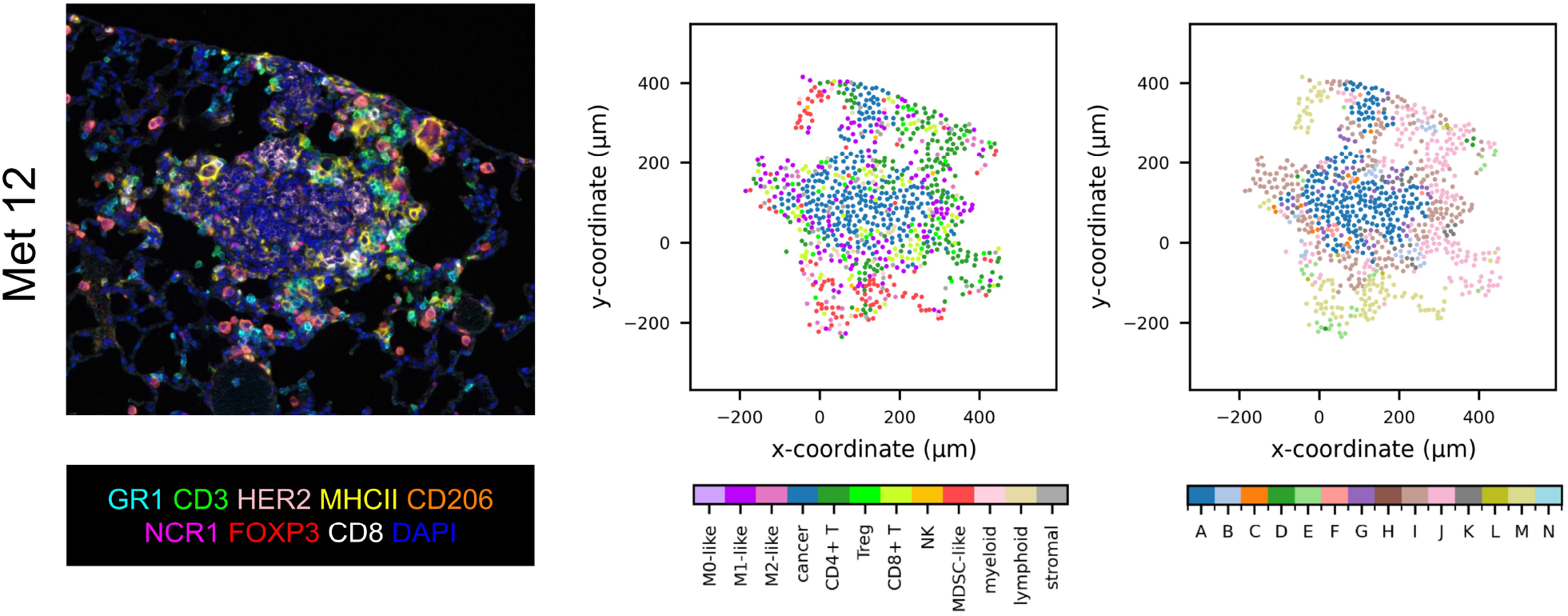
The relative spatial distribution of immune cells may explain heterogeneity in response to blockade of MDSC-like cells. mIHC image (left), cell type-labeled digitized metastasis (center), and cellular neighborhood-labeled digitized metastasis (right) for Met 12. **Alphabetical neighborhood labels:** A - uninfiltrated cancer cell region; B - CD8+ T cell region; C - MDSC-like-enriched tumor; D - M2-like macrophage region; E - M2-like macrophage/MDSC-like-enriched tumor; F - M2-like macrophage-enriched tumor; G - MDSC-like/Treg-enriched tumor; H - Treg-enriched tumor; I - T cell activation region; J - Suppressed CD8+ T infiltrated tumor; K - CD8+ infiltrated tumor; L - myeloid region; M - MDSC region; N - generalized cell zone.

## 3 Discussion

We present an ABM that captures the biological complexity of breast-to-lung metastases and reveals mechanisms of ICI resistance. Using this model, we simulated the spatiotemporal evolution of 19 lung metastases obtained from a preclinical model of metastatic breast cancer, without treatment and with dual immune checkpoint inhibitors anti-PD-1 and anti-CTLA-4. These simulations demonstrated that dual ICIs reduced growth for 37% of the metastases, consistent with survival results in the same preclinical model [24]. Analysis of the ABM’s spatial outputs showed that metastases for which dual ICIs have an effect had distinct patterns of spatial organization of tumor and immune cells at multiple scales. Furthermore, these analyses identified spatial neighborhoods involving Tregs as an important facilitator of treatment resistance to ICIs. We confirmed this mechanism by showing that Treg blockade in combination with dual ICIs led to reduced growth in 80% of metastases. Thus, we demonstrate that ABMs are well suited to uncover how the spatial organization of the TME affects responses to therapy. Furthermore, we used agent-based modeling to identify a therapeutic strategy to enhance the efficacy of ICIs in a cancer setting where ICIs have been largely ineffective.

The ABM presented here reflects the complex multicellular niche of the TME and is an expanded and more detailed model relative to other published work. Compared to prior work [17, 18, 22, 23], we included NK and MDSC-like cells in the model to more accurately capture the immune complexity of the breast-to-lung metastatic TME. Additionally, we implemented a novel method for continuous CD8+ T and NK cell exhaustion phenotypes impacted by direct and indirect interactions with other cells in the TME, allowing an investigation of a wide range of immunosuppressive mechanisms that arise in metastatic breast cancer. Combining the mechanistic detail of this model with spatial analyses of its outputs allowed us to explain which factors of a metastasis’s spatial organization influence its response to treatment, and to identify and implement alternative therapeutic strategies. We can also specifically tune TME components to the site of disease, which in this case is the lung metastatic niche.

Agent-based modeling enables a mechanistic understanding of the effect of immunotherapy on heterogeneous metastases. Model simulations showed that dual ICIs only affected cancer cell growth in a small subset of metastases, and only altered the mean irRECIST classification of a single metastasis based on normalized endpoint cancer cell population. This aligns with experimental results indicating that dual ICI in the absence of other immune modulatory therapies is not effective in mouse models of metastatic breast cancer, and is effective in a similar percentage of heavily pretreated patients with metastatic disease, in agreement with the results of the meta-analysis of dual ICIs [3]. We exploited the spatiotemporal nature of this ABM by applying emerging techniques from spatial biology to the model’s outputs to investigate which features of metastases could drive resistance to dual ICIs. We found that the spatial organization of metastases in which dual ICI treatment had an effect on the cancer cell population was noticeably distinct from metastases in which treatment did not have an effect. We showed that these distinctions were preserved across scales by analyzing the effects of pairwise cell-cell interactions, cell triplets, and cellular neighborhoods on treatment outcome.

A novel finding was that most metastases that responded to dual ICIs in simulations were dominated by cancer cell-enriched neighborhoods that were virtually devoid of immune suppressor cells, and these neighborhoods were less represented in metastases that did not respond to treatment. Specifically, the ICI-responsive metastases had fewer macrophages, MDSC-like cells, and Tregs, which our model suggests enables the existing CD8+ T cells and NK cells to drive an effective anti-tumor response. This interpretation is supported by clinical studies and analyses of patient samples that demonstrate correlations between lower frequencies of M2-like macrophages, MDSC-like cells, and Tregs in tumors and response to ICIs [27–29]. Interestingly, we found that interactions between CD8+ T cells and both M2-like and M1-like macrophages, were positively associated with treatment effect, and interactions with MDSC-like cells were associated with treatment resistance, but only to a small degree. This suggests that it is important to be able to assess the nature of the interactions to understand the effect on the anti-tumor outcome; a more simplistic analysis of the presence of a cell type will not inform response. Overall, this expanded ABM combined with spatial analyses reinforces recent work showing that the spatial organization of the TME may be prognostic of progression [30] and response to various types of treatment in breast cancer [31, 32].

A key finding from our work is that blocking Tregs from the TME has a greater effect on decreasing cancer cell populations and enhancing response to dual ICIs compared to blocking MDSC-like cells. This aligns with previous work showing that Tregs are major participants in the immunosuppressive TME of breast cancer [33], and that elevated levels of Tregs are associated with worse outcomes and resistance to ICI [34, 35]. Thus, targeting Tregs can be an impactful therapeutic strategy, as supported in previous work [36]. As a result, emerging immunotherapies that lead to Treg depletion rather than just blockade, have been studied in other cancer types with some success [37]. For example, the Fc-enhanced anti-CTLA-4 antibody botensilimab (AGEN1181) has been shown to promote intratumoral depletion of Tregs, which commonly express CTLA-4, in addition to blocking CTLA-4 interactions and binding to Fc-gamma receptors on NK cells and macrophages, leading to improved T cell activity [38]. The safety and efficacy of botensilimab in combination with the anti-PD-1 inhibitor balstilimab have been evaluated in phase 1 trials for microsatellite stable metastatic colorectal cancer and advanced and/or metastatic sarcomas. In both settings, multiple patients achieved PR per RECIST 1.1 [38, 39]. While these tumors are generally considered to be “immunologically cold,” our results indicate that a ICI combination that targets Tregs, such as botensilimab combined with balstilimab, may convert TMEs heavily infiltrated with suppressive immune cells into environments responsive to ICI.

We recognize that there are limitations to this work. Firstly, while our model includes a large immune cell repertoire, we do not have all possible immune cells that influence tumor growth and response to treatment. For example, professional antigen-presenting cells like B cells and dendritic cells are involved in T cell activation and directly participate in CTLA-4-mediated suppression caused by Tregs [40, 41]. Recent work from our group also shows that B cells are important when dual ICIs are combined with the histone deacetylase (HDAC) inhibitor entinostat [24]. Additionally, we did not include the two distinct forms of MDSCs (monocytic MDSCs and granuloctyic/polymorphonuclear MDSCs), which operate via separate immunosuppressive mechanisms, have varying effects on the TME [42, 43] and T cell activation [24]. Incorporating additional immune cells and subtypes into future versions of the ABM will enable further exploration of how spatial organization influences various therapeutic strategies.

Secondly, while we integrated experimental data throughout the model development process and initialized simulations directly from tumor imaging data, we acknowledge that we have not fully calibrated the model to experimental data. Given additional tumor images across different drug treatment regimens, future work can leverage our published approach for calibrating ABMs to imaging data [21]. Additionally, time-course data such as tumor volume can be used to further constrain the simulated model dynamics. We have also yet to account for the effect of the normal organ on the mechanics of metastatic cancer cell growth, which will be important in understanding differential responses by metastatic site.

In summary, we demonstrate that agent-based modeling can effectively capture how the spatial organization of the TME in breast-to-lung metastases contributes to resistance to immunotherapy. The model’s outputs across multiple scales highlight Tregs as a key factor limiting the effectiveness of anti-PD-1 and anti-CTLA-4 combination therapy. We further applied the model to evaluate Treg blockade as a potential therapy concurrent with dual ICIs, and showed that removing Tregs increases treatment efficacy across the cohort of simulated metastases. Thus, this model points to a potential method for converting highly immunosuppressive TMEs into environments responsive to ICIs, demonstrating the utility of ABMs for rapidly testing therapeutic strategies in solid tumors.

## 4 Methods

### 4.1 *In vivo* studies

#### 4.1.1 Cell lines

NT2.5-LM is a rat HER2/neu+ breast cancer cell line derived from NT2.5 (RRID:CVCL JA60) that spontaneously metastasizes to the lungs of NeuN mice with high penetrance. The NT2.5-LM cell line was obtained from the lab of Elizabeth Jaffee at Johns Hopkins School of Medicine. NT2.5-LM was cultured in RPMI-1640 base media (Gibco, cat. #11875-093) supplemented with 20% fetal bovine serum (Gemini, cat. #100-106), 1.2% HEPES (Gibco, cat. #15630-080), 1% l-glutamine (Gibco, cat. #25030-081), 1% non-essential amino acids (Gibco, cat. % 11140-050), 1% sodium pyruvate (Sigma, cat. #S8636), 0.5% penicillin/streptomycin (Gibco, cat. #15140-122), and 0.2% insulin (NovoLog, cat. #NDC 0169-7501-11) at 37 °C and 5% carbon dioxide. NT2.5-LM cells were submitted to Charles River Research Animal Diagnostic Services to test for contamination with mycoplasma and other pathogens, with the last test performed on Oct. 25, 2025. NT2.5-LM cells were submitted to ATCC for authentication but could not be successfully authenticated due to NT2.5-LM not appearing on the ATCC database. NT2.5-LM cells were thawed and kept in culture for a minimum of five days and passaged twice before injection into mice.

#### 4.1.2 Mice experiments

Female NeuN mice (Jackson Labs strain #002376) were bred in-house in the animal housing facility at the University of Southern California. Mice were housed under pathogen-free conditions and handled according to protocol 21770 approved by the USC Institutional Animal Care and Use Committee (IACUC), following guidelines from the American Association for Laboratory Animal Committee policies.

NeuN mice were injected with 100,000 NT2.5-LM cells in the 2nd mammary fat pad. Tumors were surgically resected on day 7 post-inoculation for all samples except those collected on day 7. Lungs were collected on days 7, 10, 16, 22, 28, 33, 35, and 122 post-inoculation. Lungs were inflated and fixed in 10% neutral-buffered formalin and embedded in paraffin before sectioning for IHC analysis. The specific metastases used in this study, their mice of origin, and the corresponding harvest day (post-inoculation), are shown in **Sup. Table S1**.

#### 4.1.3 Multispectral IHC imaging

Through our collaboration with the Human Immune Monitoring Shared Resource (HIMSR) at the University of Colorado School of Medicine, we performed multispectral imaging using the PhenoImager HT instrument (Akoya Biosciences). To quantify levels of fluorescence, formalin-fixed paraffin-embedded tissue sections were stained consecutively with specific primary antibodies for HER2, CD3, CD8, FOXP3, GR1, MHCII, CD206, and NCR1. Briefly, the slides were deparaffinized, heat treated in antigen retrieval buffer, blocked, and incubated with primary antibodies, followed by horseradish peroxidase (HRP)-conjugated secondary antibody polymer, and HRP-reactive OPAL fluorescent reagents. The slides were stripped between each stain with heat treatment in antigen retrieval buffer. Whole slide scans were collected with PhenoImager HT v2.0.0 software using the 20x objective with a 0.5 micron resolution. Regions of interest were selected and rescanned using the 20x objective and the multispectral imaging cube. Spectral references and unstained control images were measured and inForm software v3.0 was used to create a multispectral library reference. The multi-spectral images (.im3 files) were spectrally unmixed, analyzed with tissue segmentation, cell segmentation, and phenotyping using inForm software v3.0 (Akoya Biosciences), and data were compiled and summarized using PhenoptrReports (Akoya Biosciences).

### 4.2 *In silico* agent-based model of the tumor microenvironment

Here, we provide a brief overview of the agent-based model, which is based on our prior work [22, 44], and detail the primary changes we have made for the present study. A thorough description of the model is contained in the Supplementary Methods, and its implementation in C++ is available on GitHub (see “Code and data availability” section below). To account for stochastic effects inherent to agent-based modeling, we simulate 25 replicates per treatment condition for each metastasis. For Metastases 6 and 14, we simulate an additional 75 replicates per treatment condition to better explore the divergent behavior predicted by the model.

Our model records the spatial organization of the simulated metastasis every hour for a simulation window of 25 days (600 hours). In addition to recording the spatial location of every cell, the model also captures the cell’s current state, its PD-1 or PD-L1 expression, and its cell cycle length (for cells allowed to proliferate). For CD8+ T cells and NK cells, the model also records details about the cell’s exhaustion phenotype.

#### 4.2.1 Cell types

The ABM includes cancer cells, CD8+ T cells, NK cells, macrophages, CD4+ T cells, T regulatory cells, and MDSC-like cells. It also includes three types of “generalized” cells that do not have biological behavior but are present to maintain tissue structure: myeloid cells, lymphoid cells, and stromal cells. As in our previous work, macrophages can polarize into the anti-tumor M1-like state or the pro-tumor M2-like state, and they enter simulation in the näıve M0-like state. Additionally, CD4+ T cells can irreversibly differentiate into T regulatory cells. **Fig. 1B** provides a summary of the non-generalized cell types, their interactions with each other, and the cell processes they undergo.

#### 4.2.2 Physics

The model uses the overlapping spheres method to describe cells and their physical interactions with one another. For each cell, we track its center position and its radius. In brief, cells migrate through the simulation environment using random walks with step lengths determined by their migration speed. If two cells overlap each other, then a repulsive force pushes them apart. Additionally, there are cohesive forces between cancer cells. Collectively, these forces also cause cells to move. The exact equations describing these forces are available in the **Supplementary Methods**.

#### 4.2.3 Cell-cell interactions

The ABM contains two types of cell-cell interactions: direct interactions, which require physical contact between two cells to occur, and indirect interactions. The two forms of direct interactions, which involve either effector cells killing cancer cells or PD-1/PD-L1 mediated exhaustion of effector cells, are described in detail below. Indirect cell interactions involving only macrophages, CD4+ T cells, T regulatory cells, CD8+ T cells, and cancer cells have been previously described in [22]. Below, we discuss indirect interactions that involve the two cell types added in this iteration of the model compared to our prior work, MDSC-like cells and NK cells.

##### 4.2.3.1 Indirect interactions

In the TME, diffisible factors such as cytokines mediate indirect interactions between cells. We do not explicitly model diffusible factors and instead use the cellular influence framework we previously developed [22]. Briefly, this framework treats each cell as the source of a field that decays exponentially, and the value of this field a defined distance away from the cell center gives the probability that an indirect interaction occurs due to the influence of a nearby cell. Thus, we can quickly examine all cells of a specific type within the neighborhood of a cell of interest and evaluate the cell type’s total influence by calculating the probability that no event occurs due to that cell type and taking the complement. The explicit form of these equations is available in the Supplementary Methods.

##### 4.2.3.2 MDSC-like cell interactions

Myeloid-derived suppressor-like (MDSC-like) cells are a class of immature myeloid cells that can potentially differentiate into monocytes and macrophages, and have a range of immunosuppressive functions in the TME. In our model, MDSC-like cells are involved in the induction of PD-1 onto CD8+ T cells [45] and NK cells [46], and also promote the exhaustion of CD8+ T cells and NK cells [42, 47–53]. Additionally, MDSC-like cells promote macrophage polarization into the M2-like state [54], and the differentiation of CD4+ T cells into T regulatory cells [50].

As described below, the expression of PD-L1 on the surface of MDSC-like cells can be induced by CD8+ T cells, CD4+ T cells, and NK cells, which is shown to occur through IFN-*γ* secretion [55].

##### 4.2.3.3 NK cell interactions

Natural killer (NK) cells are a type of innate lymphoid cell which can engage in the killing of cancer cells. In our model, NK cells behave similarly to CD8+ T cells in that they have an exhaustion phenotype, form immune synapses with cancer cells, and can kill cancer cells; however, these cells do not proliferate and are not affected by the anti-CTLA-4 drug. As described below, the expression of PD-1 on the surface of NK cells can be induced by cancer cells [56], Tregs [57], M2-like macrophages, and MDSC-like cells [46]. Additionally, NK cells are exhausted directly via PD-1/PD-L1 interactions, and indirectly via Tregs, M2-like macrophages, and MDSC-like cells [53, 57]. Conversely, the presence of M1-like macrophages [58, 59] and CD4+ T cells [60] have a restorative effect on the exhaustion phenotype of NK cells, making NK cells more active.

NK cells are also involved in the induction of PD-1 onto cancer cells [61] and the protumor immune cell types (M2-like macrophages, MDSC-like cells, and Tregs). NK cells can influence macrophage polarization into the M1-like state [59, 62] via IFN-*γ* secretion [63]. Finally, NK cells can indirectly have an anti-exhaustive effect on CD8+ T cells [64].

##### 4.2.3.4 Immune synapses

When CD8+ T cells and NK cells make contact with and recognize their target (cancer cells), they form an immune synapse, which physically connects them to the target cell. These synapses allow cytotoxic cells to push lytic granules into the target cell to promote apoptotic signaling [65–67]. We model this interaction by introducing an additional spring force that pulls the cells toward each other. We set the resting length of the spring (*l*_rest_)to be slightly smaller than the sum of the two cells’ radii, such that when the spring force is zero, the two cells overlap. The strength of this force (given by the spring constant *k*_syn_) has been tuned so the immune synapse force dominates the repulsive forces caused by the overlap of the two cells, and is given by:

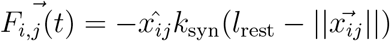

In order for a CD8+ T cell or NK cell to kill a cancer cell, there must be an immune synapse connecting the two cells together. These cells can form up to two immune synapses simultaneously, and each synapse remains active for one hour after initial contact.

#### 4.2.4 CD8+ T cell and NK cell exhaustion phenotypes

The behavior of effector cells in our model (CD8+ T cells and NK cells) is governed by five cell-intrinsic parameters: killing probability, migration speed, migration bias, life span, and cell cycle length. Each of these parameters can be decreased by the cellular program known as “exhaustion,” wherein the effector cells lose cytotoxic capacity, upregulate inhibitor receptors, and undergo a number of other physical and behavioral changes [53, 68, 69]. We model these changes by changing these cell-intrinsic parameters; thus, we describe a continuum of exhaustion states rather than a subset of discrete states. Note that NK cells do not proliferate in our model, so we do not track changes in their cell cycle length.

The rate of exhaustion of a specific cell parameter is controlled by a scaling parameter *k_d_*, and the amount of exhaustion that a cell experiences is a function of either indirect cell influences or the strength of a direct PD-1/PD-L1 interaction. If the strength of this interaction at time *t* is described by the function *D*(*t*), then the change in a cell-intrinsic parameter *p* can be described as:

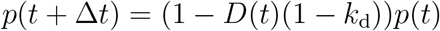

In the case where cells experience the maximum amount of exhaustion (e.g., *D*(*t*) = 1 for all *t*), then the equation above approximates exponential decay in the parameter over time. Cell cycle length and lifespan are tracked as their inverse (e.g., as the probability of division and the probability of death). Because the probability of death increases with exhaustion, unlike the other parameters that decrease with exhaustion, it is adjusted in a slightly different manner:

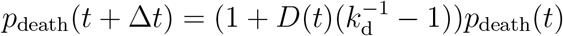

For indirect interactions, the value of *D*(*t*) for a specific parameter is equal to the combined influence of the cell types that affect that parameter (see the **Supplementary Methods**). Additionally, the values of k_d_ associated with each exhaustion property and cytotoxic cell type and their derivations are given in the **Supplementary Methods**).

##### 4.2.4.1 PD-1/PD-L1 exhaustion and anti-PD-1 drug

The immune checkpoint molecule PD-1 can be expressed on CD8+ T cells and NK cells, and its ligand PD-L1 can be expressed on the pro-tumor cells present in our model (cancer cells, M2-like macrophages, MDSC-like cells, and T regulatory cells). When an effector cell is in direct contact with a PD-L1-bearing cell, interactions between the PD-1 and PD-L1 molecules can lead to cell exhaustion.

The expression of PD-1 on effector cells is controlled by the combined influence *D*_+_ of cancer cells, M2-like macrophages, MDSC-like cells, and Tregs. If *D*_+_ is greater than a given threshold, *P*_th,PD-1_, then PD-1 is expressed at a rate proportional to the difference between *D*_+_ and *P*_th,PD-1_. If *D*_+_ is less than the threshold, the PD-1 decays at a constant rate:

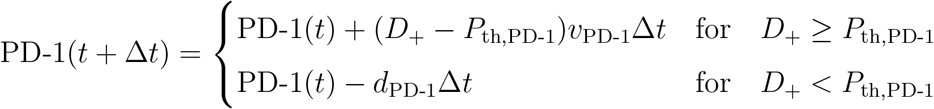

The expression of PD-L1 on the pro-tumor immune cells and cancer cells is controlled by the combined influence *D*_+_ of CD8+ T cells, CD4+ T cells, and NK cells. The equation follows the same form as the PD-1 expression equation using the induction threshold *^P^*_th,PD_*_−_*_L1_:

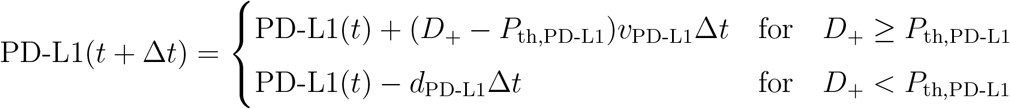

We additionally simulate an anti-PD-1 drug that can sequester PD-1 molecules from participating in these interactions. If the anti-PD-1 drug is present, we assume that the drug and PD-1 quickly reach an equilibrium. Thus, the amount of PD-1 available to interact with PD-L1 molecules can be given by:

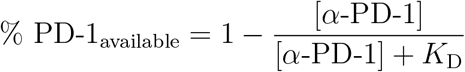

We assume that the maximal amount of binding occurs between available PD-1 and PD-L1. We can then determine the exhaustive influence *D*(*t*) exerted by this interaction by comparing the amount of PD-1 bound to the maximum amount of PD-1 expression allowed in the model. We scale this effect such that when a cell expresses half the maximum amount of PD-1 and these molecules are fully bound to PD-L1 molecules, *D*(*t*) = 1:

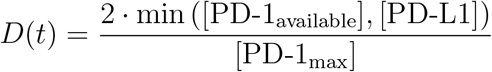

If an effector cell is “severely exhausted,” meaning that its killing probability, migration speed, and lifespan are 25% of baseline, we assume that the cell is resistant to the drug and that the amount of PD-1 available is equal to the amount of PD-1 expressed, regardless of the drug concentration. The anti-PD-1 drug is administered at intervals determined by the experimental treatment schedule shown in **Fig. 1C**. We assume the concentration of the drug is the same across the simulation domain, and decays according to first-order kinetics with a decay coefficient *k*_decay_ = 1.93 *×* 10*^−^*^3^.

##### 4.2.4.2 CTLA-4 mediated exhaustion and anti-CTLA-4 drug

The expression of CTLA-4 on CD8+ T and T regulatory cells can also contribute to the exhaustion of CD8+ T cells by inhibiting the pathways needed to maintain effector cell activity. Because NK cells do not participate in the CTLA-4 pathway, they are not affected by CTLA-4 or the anti-CTLA-4 drug. In our model, this pathway specifically affects the indirect interaction update to the cell cycle length of the CD8+ T cell.

Specifically, the influence of CD4+ T cells and NK cells can decrease the cell cycle length (thus increasing proliferation), while the influence of M2-like macrophages and MDSC-like cells increases the cell cycle length. The positive and negative influences can thus be expressed as:

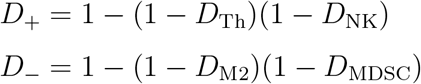

T regulatory cells, which can express CTLA-4, modulate the value of the positive influence *D*_+_. The strength of this modulation, *η*, depends on the concentration of anti-CTLA-4 drug in the environment, which we model using a Hill equation. The value for the activation constant, *k*_act_, is increased to be 1.1 for cells with a cell cycle length shorter than baseline; for other cells *k*_act_ = 1.

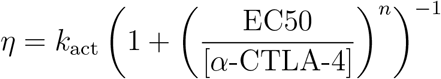

Thus, from the form of the exhaustion equation above, we have:

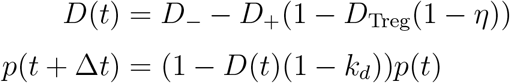

As with the anti-PD-1 drug, the anti-CTLA-4 drug is administered at intervals determined by the experimental treatment schedule shown in **Fig. 1C**. We also assume the concentration of the drug is the same across the simulation domain, and decays according to first-order kinetics with a decay coefficient *k*_decay_ = 8.53 *×* 10*^−^*^4^.

#### 4.2.5 Preparation of initial conditions from imaging data

Eighty potential metastases were manually identified from mIHC images and annotated. The associated cells were identified and retrieved from the processed imaging data using a custom MATLAB script which is available at the GitHub repository for this paper.

##### 4.2.5.1 Cell type assignment

Binary protein expression profiles and their associated fluorescence values were obtained from imaging as described above. Briefly, cells were categorized as positive for one of the eight markers in the IHC panel if the expression level surpassed an algorithmically-determined threshold. Cells were then binned into different categories based on their expression of each marker. Using binary protein expression, cells were labeled as MDSC-like cells, M1-like macrophages, M2-like macrophages, NK cells, Tregs, CD8+ T cells, and CD4+ T cells if they expressed markers shown to identify these cells types (see **Supplementary Table 1** for all combinations of cell markers used to assign each cell type). Since cancer cells are genetically unstable and can upregulate genes not typically observed in the cell of origin, cancer cells were defined as HER2+, along with any combination of the remaining markers, except for two NK cell combinations noted in the table. If a cell could not be labeled as a cancer cell or immune cell type using this method, it was assigned an intermediate “unknown” cell type.

Cells assigned an unknown cell type based on binary protein expression and marker combinations given in the look-up table were assigned a cell type using a random forest (RF) model. The random forest model was trained using the cells successfully labeled using the look-up table. The input features for the random forest were the continuous fluorescence values from mIHC. Unlabeled cells were assigned either a defined cell type or a generalized cell type based on the class probabilities predicted by the RF model (**Supplementary Fig. S1**). Briefly, if the highest class probability is sufficiently large relative to other classes, then the cell is assigned into that cell type. However, if this is not the case, then we examine the relative (summed) probabilities of the immune lineages to determine whether a cell should be assigned into the generalized types of “myeloid” or “lymphoid,” or if it should be assigned to the non-immune “stromal” class. The RF model is available in the GitHub repository for this paper.

##### 4.2.5.2 Rescaling

Cells in the model have fixed, pre-defined radii based on literature evidence, and the cell physics model we use is tuned to cells of these sizes. To ensure that metastases are not overly-compressed at early time points, which would lead to the non-biological behavior of a large, rapid spreading of the cells to compensate for tight packing, we scale the coordinate axes of the mIHC data equally in both directions such that cells, on average, do not overlap at time zero. We calculate this scale factor by computing the estimated actual density (*ρ_a_*) and estimated required density (*ρ_t_*) of the metastasis. To calculate *ρ_a_*, we first estimate the area of the metastasis by computing the alpha-shape of the cell centers with a sufficient radius parameter to enclose all cells within the metastasis with a single contour. We compute this alpha-shape using the alphashape Python package [70]. The boundary of the alpha-shape approximates the actual boundary of the metastasis, and thus its enclosed area is approximately the actual area of the metastasis. Thus, *ρ_a_* is simply the number of cells in the metastasis divided by the enclosed area. To calculate *ρ_t_*, we consider the total area that would be covered by all cells in the metastasis given our predefined cell radii. However, because cells (which we approximate as circles) do not fully cover the plane, we scale each area by a constant 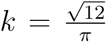 to account for the area of the plane not covered by the cells. Thus, for a metastasis with n cells, where the ith cell has radius r*_i_*:

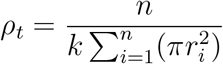

The scale factor, s, is the square root of the ratio between the two values:

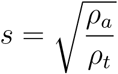

To scale the mIHC coordinate system to the simulation coordinate system, we simply multiply the x-y coordinates that define each cell center by s.

##### 4.2.5.3 Selection of initial conditions to simulate

After all cells had been assigned a cell type, either by look-up table or random forest model, metastases were screened based on their proportion of generalized cells, their size, the proportion of cancer cells, and their proportion of CD8+ T cells. Of the initial 80 metastases retrieved from the imaging data, we rejected those that contained:

- *>* 40% generalized cells (n = 35),
- *>* 25% CD8+ T cells AND *<*5 % cancer cells (n = 5),
- *<* 3% cancer cells (n = 20),
- *<* 50 cells total (n = 1).

After filtering via these criteria, we proceeded to simulate 19 metastases.

#### 4.2.6 Sensitivity Analysis

We performed sensitivity analyses, individually varying five parameters related to CD8+ T cells in the model: the CD8+ T cell recruitment rate, the k*_d_* value of p(kill), the k*_d_* value for cell cycle length, the k*_d_* value for lifespan, and the base value of p(kill). For these analyses, we selected four metastases (Mets 0, 7, 30, and 34) that had different initial proportions of CD8+ T cells, and ran 10 replicates of each metastasis for each parameter value we evaluated.

#### 4.2.7 Targeted blockade of specific cell types

We simulated blockade of MDSC-like cells and Tregs. Blockade of MDSC-like cells was performed by preventing their recruitment to the metastasis. Blockade of Tregs was performed by preventing CD4+ T cells from differentiating into Tregs. No adjustments were made to the average lifespan of these cells in simulation. Cell blockade was simulated without dual ICIs, and we performed 25 replicates. Additionally, the initial conditions of each metastasis were altered so any instance of the blocked cell was converted to the generalized cell of the proper immune lineage.

### 4.3 Data analysis

#### 4.3.1 Quantifying treatment effect

For each replicate, we calculate the normalized number of cancer cells at the simulation endpoint (600 hours) by dividing the number of cancer cells at this time by the number of cancer cells present in the initial conditions. We then use a two-sided student’s *t*-test to determine whether treatment (dual ICIs, blockade of MDSC-like cells, Treg blockade, or combinations of these treatments) has an impact on final outcomes. We perform this statistical test for all metastases excluding Mets 6 and 14, which have non-normal outcome distributions. For these metastases, we instead use the Mann-Whitney *U* test.

Additionally, we classify each replicate based on the irRECIST criteria, as either complete response (CR, complete elimination of cancer cells), partial response (PR, reduction in cancer population by > 30%), stable disase (SD, change in cancer population between -30% and +20%), or progressive disease (PD, increase in cancer population by > 20%).

#### 4.3.2 Analysis of treatment on CD8+ T cell exhaustion phenotypes

For every CD8+ T cell in a simulation replicate, we retrieve and sum its exhaustion phenotype features (i.e., migration speed, migration bias, probability of killing, cell cycle length, and lifespan; these features are normalized relative to baseline) over its lifetime to produce an “exhaustion trajectory” and a single score that approximates the cell’s lifetime activity. We then consider the distribution of scores associated with the probability of killing for these cells.

Qualitative analysis shows that these scores fall into two distributions: an exponentially-distributed population with low values and a normally-distributed population with higher values. We call these the “rapidly-exhausting” and “non-rapidly exhausting” populations. We select a threshold of *p*(*kill*) score *>* 5 to separate these populations for all metastases.

To evaluate the effect of the dual immune checkpoint inhibition therapy, we evaluate the mean and median scores for the non-rapidly exhausting population for each replicate, and use either a two-sided student’s *t*-test or Mann-Whitney *U* test to determine whether the non-treatment and treatment groups are significantly different.

#### 4.3.3 Pairwise cell co-localization analysis

Pairwise cell-cell co-localization is calculated using the “co occurence matrix” function in the TACCO package [71], with the maximum distance set to 30 *µ*m and the “score key” parameter set to “z.” This distance is chosen to be three times the cancer cell radius (i.e., this region will completely enclose a cancer cell exactly tangent to the center of a cancer cell of interest). This method adapts the procedure in [30], which compares the number of neighbors within the distance threshold for a given tissue section to the expected number of neighbors if the cell type labels for the tissue section are randomly permuted. The Z-score is calculated relative to the distribution generated by 1000 random permutations; thus, a Z-score of zero indicates that the co-localization metric calculated is equal to what would be expected for a tissue segment with no structure (random cell type labels). The matrix contains the Z-score evaluated for every pairwise combination of cell types excluding generalized cells and including a cell type’s co-localization score with itself. If a cell type is not present in a specific simulation output, we set the z-scores for all pairwise combinations associated with that cell equal to zero such that every matrix is the same shape.

For each simulated non-treatment replicate, we calculate this co-localization matrix every 24 hours. We then reshape each matrix into a vector. To visualize the similarity between these vectors, we use a UMAP to project each vector into 2D space. For the UMAP, we used the Python implementation [72] with n neighbors = 10, min dist = 0.5, and random state=42. Since the co-localization matrix is symmetric, we only consider the upper triangular entries when converting these matrices into vectors for input into UMAP.

#### 4.3.4 Interaction ratios for CD8+ T-cancer pairs

For each simulation output, we construct a Delaunay triangulation of the spatial organization to determine possible direct interactions and strong indirect interactions. We use the SciPy implementation of Qhull to calculate the Delaunay triangulation [73], and prune edges in the triangulation that are greater than 20 *µ*m (i.e., two cancer cell radii). Using this graph, we determine the subset of CD8+ T cells that have a cancer cell as a neighbor, and thus participate in a “CD8+ T–cancer dyad.” Subsequently, for every immune cell type present in the model, we determine the number of CD8+ T cells in this subset that have a neighbor of the specified immune cell type, and thus participate in a cancer–CD8+ T cell–immune cell triplet (see **Fig. 5A** for an example of a cancer-CD8+ T cell-MDSC-like cell triplet). We do not calculate such triplets for M0-like macrophages or the generalized cells, since those cells do not participate in cell-cell interactions. Additionally, if a CD8+ T cell of interest has multiple immune neighbors, the same cancer-CD8+ T cell dyad will be counted as part of multiple triplets. Furthermore, if a cancer cell has multiple CD8+ T cell neighbors in the Delanunay triangulation, it will participate in multiple cancer-CD8+ T cell dyads.

We then define the “interaction ratio” (IR) for a given immune cell as the ratio of CD8+ T cells in this triplet divided by the number of CD8+ T cells in CD8+ T – cancer dyads:

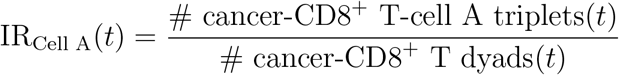

For every metastasis, we calculate each immune cell’s IR for each replicate at every hour simulated. We can additionally define a “cancer vulnerability” score by counting the number of cancer cells with a CD8+ T cell neighbor and dividing this by the total number of cancer cells:

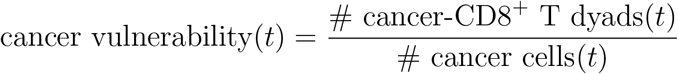

#### 4.3.5 Quantifying relative impact of IRs on treatment response

For each replicate, we convert the time series IR data into a single score per immune cell by calculating the area under the IR curve for each immune cell for the first 480 hours of simulation. We convert the time series cancer vulnerability data into a single score using the same procedure. Thus, we produce a set of eight scores associated with each simulated replicate.

To identify whether certain cancer–CD8+ T–immune cell triplets are associated with a metastasis experiencing a treatment effect, we trained a logistic regression using the eight scores as input features and whether or not a metastasis experienced treatment effect as the output feature. Prior to training, each feature was min-max normalized across the dataset.

To determine if pairwise combinations of IR scores and/or cancer vulnerability scores would be better inputs for the logistic regression, we developed two additional regression models that contained multiplicative pairwise combinations of features (we do not include the product of a feature with itself) and identified the optimal integration window via 5-fold cross validation as before. After determining the ideal window, we trained the models on the full data set to investigate the regression coefficients.

#### 4.3.6 Cellular neighborhood analysis

We define cellular neighborhoods based on the spatial outputs of simulation as described in [74]. In brief, for each cell, we identify its cell type and the cell types of its nine nearest neighbors, then construct a vector that indicates the number of times each cell type appears in this set of ten cells. We construct these vectors for all cells in the first 48 hours of all no-treatment replicates. We then cluster these vectors together using minibatch k-Means with k = 30. To label each cluster with a semantically-meaningful name, we identify the centroid position for each cluster and also evaluate the log-fold change of each cell type for the centroid position relative to the average cell in the data set. The assignment of cluster labels is available in **Supplementary Table 2**; additionally, cluster definitions are shown in **Sup. Fig. S7**. By manually labeling clusters, we reduce the number of unique clusters from 30 to 14. We evaluated alternative clusterings with k = 15 and k = 40, and found that increasing the number of clusters would lead to most cells being assigned to the same manually labeled neighborhood. However, decreasing the number of clusters would lead to clusters that contained multiple manually labeled neighborhoods, leading to data loss. Thus, we used k = 30 for the analysis.

For each metastasis, we calculate the fraction of cells in the initial condition that are assigned to each neighborhood cluster. We then hierarchically cluster these vectors together using the default clustering method built in to the “clustermap” function of Seaborn (the linkage method defaults to “average,” which is equivalent to the UPGMA algorithm, and we use the default Euclidean distance metric) [75]. Additionally, we repeat this clustering using the fraction of cells in the initial condition that are assigned to each cell type, to determine whether spatial information in addition to cell identity predicts whether simulated treatment will have an effect on the growth of the cancer cell population.

## 4.4 Code and data availability

All code is available at the project’s GitHub repository, including the C++ implementation of the agent-based model and the Python code necessary for conducting analyses. The repository includes a complete list of dependencies needed to run both the C++ and Python code that produces the results. The C++ implementation of the model must be compiled and run on Unix-based architecture. Large sections of this code are parallelized to take advantage of high core-count CPU architecture, and thus OpenMP is required to run the model.

The repository also contains a Jupyter notebook that generates all figures in this paper from simulation output data. The raw simulation outputs are available upon request. We also include the slurm and input files used to specify these simulations. Pre-processed data for some computationally-expensive analyses are provided, but can also be generated on-the-fly using the Jupyter notebook. We will also provide the proteomics data generated for this study upon request.

## 5 Supplementary Material

**Supplementary Table S1:** Mice of origin and harvest day for the 19 simulated metastases.

| Metastasis | Mouse of Origin | Day Harvested |
| --- | --- | --- |
| Met 32 | Mouse 10 | 22 |
| Met 6 | Mouse 16 | 122 |
| Met 40 | Mouse 17 | 33 |
| Met 12 | Mouse 19 | 7 |
| Met 49 |  |  |
| Met 22 | Mouse 27 | 35 |
| Met 45 |  |  |
| Met 25 | Mouse 28 | 28 |
| Met 8 | Mouse 29 | 28 |
| Met 23 |  |  |
| Met 28 |  |  |
| Met 30 |  |  |
| Met 0 | Mouse 31 | 28 |
| Met 7 |  |  |
| Met 13 |  |  |
| Met 14 |  |  |
| Met 17 |  |  |
| Met 34 |  |  |
| Met 43 |  |  |

**Supplementary Table S2:** Protein expression phenotypes used to identify cell types.

| Cell type | Accepted phenotypes |
| --- | --- |
| Cancer | HER2+ / CD3+HER2+ / MHC-II+HER2+ /<br>FOXP3+HER2+ / CD3+FOXP3+HER2+ /<br>CD3+HER2+MHC-II+ / CD3+CD8+HER2+ /<br>CD3+CD8+HER2+MHC-II+ / CD8+HER2+MHC-II+ /<br>CD8+HER2+ / CD3+CD8+HER2+FOXP3+ /<br>CD3+FOXP3+HER2+MHC-II+ / FOXP3+HER2+MHC-II+ /<br>CD3+CD8+FOXP3+HER2+MHC-II+ /<br>CD8+HER2+NCR1+ / CD8+FOXP3+HER2+ /<br>CD206+FOXP3+HER2+ / CD206+CD3+FOXP3+HER2+ /<br>CD3+CD8+HER2+NCR1+ / GR1+HER2+ /<br>CD206+CD3+HER2+ / CD206+CD3+CD8+FOXP3+HER2+ /<br>CD3+GR1+HER2+ / CD3+FOXP3+HER2+NCR1+ /<br>CD206+FOXP3+HER2+MHC-II+ / CD206+CD8+HER2+ /<br>CD206+HER2+ / CD3+CD8+GR1+HER2+ /<br>CD206+CD3+CD8+HER2+ /<br>CD8+FOXP3+HER2+MHC-II+ / GR1+HER2+MHC-II+ |
| MDSC | GR1+ |
| M1-like macrophage | MHC-II+ / GR1+MHC-II+ / FOXP3+GR1+MHC-II+ /<br>CD8+GR1+MHC-II+ / CD3+CD8+GR1+MHC-II+ /<br>CD3+GR1+MHC-II+ |
| M2-like macrophage | CD206+ / CD206+FOXP3+ / CD206+FOXP3+MHC-II+ /<br>CD206+MHC-II+ / CD206+CD8+MHC-II+ /<br>CD206+CD8+MHC-II+FOXP3+ /<br>CD206+CD3+CD8+MHC-II+ / CD206+FOXP3+GR1+ /<br>CD206+GR1+ / CD206+CD3+FOXP3+MHC-II+ /<br>CD206+CD3+FOXP3+NCR1+ / CD206+GR1+MHC-II+ |
| NK cell | NCR1+ / NCR1+ HER2+ / CD3+ HER2+ NCR1+ / CD3+<br>NCR1+ |
| CD8+ T cell | CD8+ / CD3+ CD8+ / CD8+ GR1+ / CD3+ CD8+ GR1+ |
| CD4+ T cell | CD3+ / CD3+ GR1+ |
| Tregs | FOXP3+ / CD3+ FOXP3+ / CD3+ FOXP3+ GR1+ |

**Supplementary Table S3:** Mapping of numerical clusters to semantic labels.

| **Numeric Cluster** | **Semantic Cluster Name** | **Alphabetical Label** |
| --- | --- | --- |
| 3 | tumor | A |
| 6 |  |  |
| 15 |  |  |
| 21 |  |  |
| 23 |  |  |
| 4 | CD8+ T cell region | B |
| 7 |  |  |
| 11 |  |  |
| 24 |  |  |
| 10 | MDSC-like enriched tumor | C |
| 16 |  |  |
| 29 |  |  |
| 8 | M2 region | D |
| 9 | M2-like/MDSC-like enriched tumor | E |
| 18 |  |  |
| 2 | M2-like enriched tumor | F |
| 28 | MDSC-like/Treg enriched tumor | G |
| 27 | Treg enriched tumor | H |
| 5 | T cell activation region | I |
| 12 | suppressed CD8+ T cell infiltrated tumor | J |
| 20 |  |  |
| 0 | CD8+ T cell infiltrated tumor | K |
| 19 | myeloid region | L |
| 26 |  |  |
| 17 | MDSC-like region | M |
| 22 |  |  |
| 1 | Generalized cell region | N |
| 13 |  |  |
| 14 |  |  |
| 25 |  |  |

**Supplementary Figure S1:**
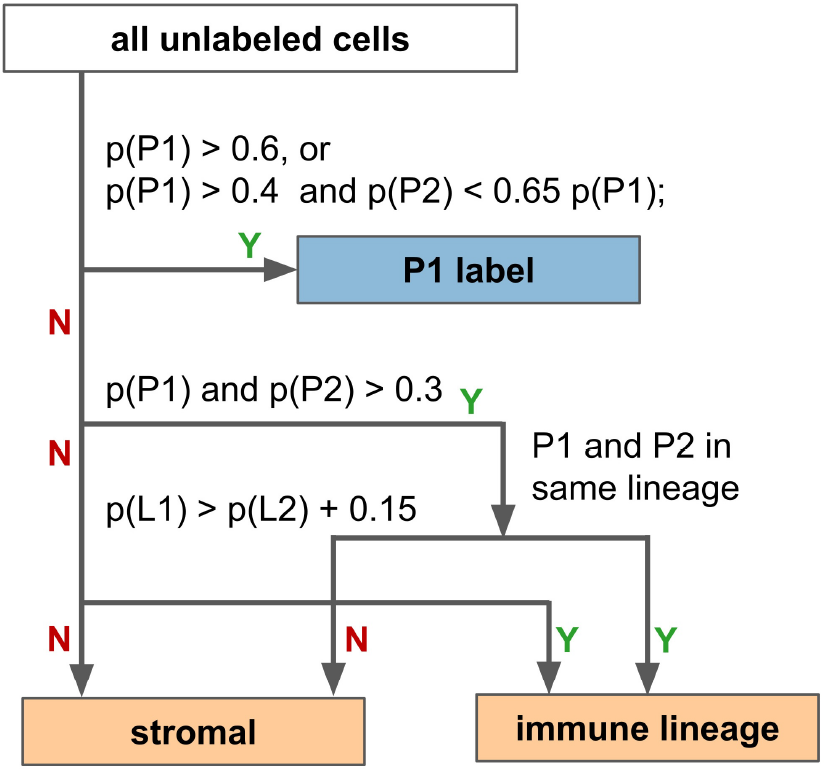
Decision tree for assigning cell types based on the class probabilities vector (output) from the random forest model. P1 indicates the cell type with the highest probability assigned by the model, p(P1). Likewise, P2 indicates the cell type with the second-highest probability, p(P2). L1 indicates the immune cell lineage with the highest total probability (summed across member cell types), p(L1), and L2 indicates the immune cell lineage with the second-highest probability, p(L2). The myeloid lineage includes MDSC-like cells, M1-like cells, and M2-like cells. The lymphoid lineage includes CD8+ T cells, CD4+ T cells, NK cells, and Tregs.

**Supplementary Figure S2:**
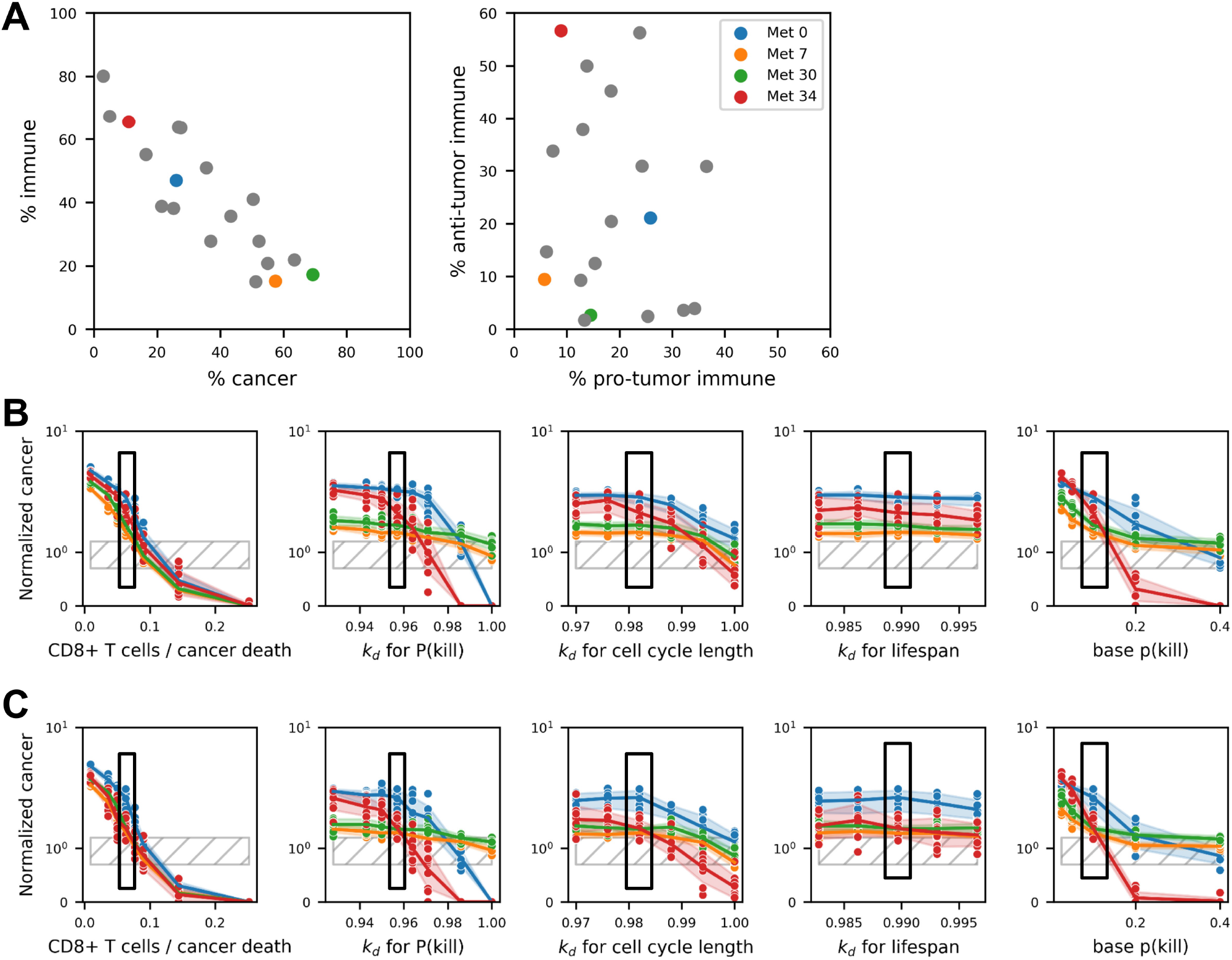
Parameter sensitivity analysis for parameters related to CD8+ T cell behavior confirms that selected parameters produce outcomes observed in experimental studies. **A.** Scatter plots showing the heterogeneity in cellular and immune-specific composition across the entire metastasis set and metastases selected for sensitivity analysis (in color). **B-C.** Sensitivity analysis was conducted for CD8+ T cell recruitment, exhaustion rate (*k_d_* values) for the probability of killing, cell cycle length, and lifespan, and for the baseline probability of killing (separate panels, left to right) in Mets 0, 7, 30, and 34. Each plot displays the normalized final cancer population (at 600 hours) as a function of the parameter value for four metastases with varying CD8+ T cell initial proportions, either with no treatment (B) or with dual ICIs (C). The parameter value used in model simulations is highlighted in a black box.

**Supplementary Figure S3:**
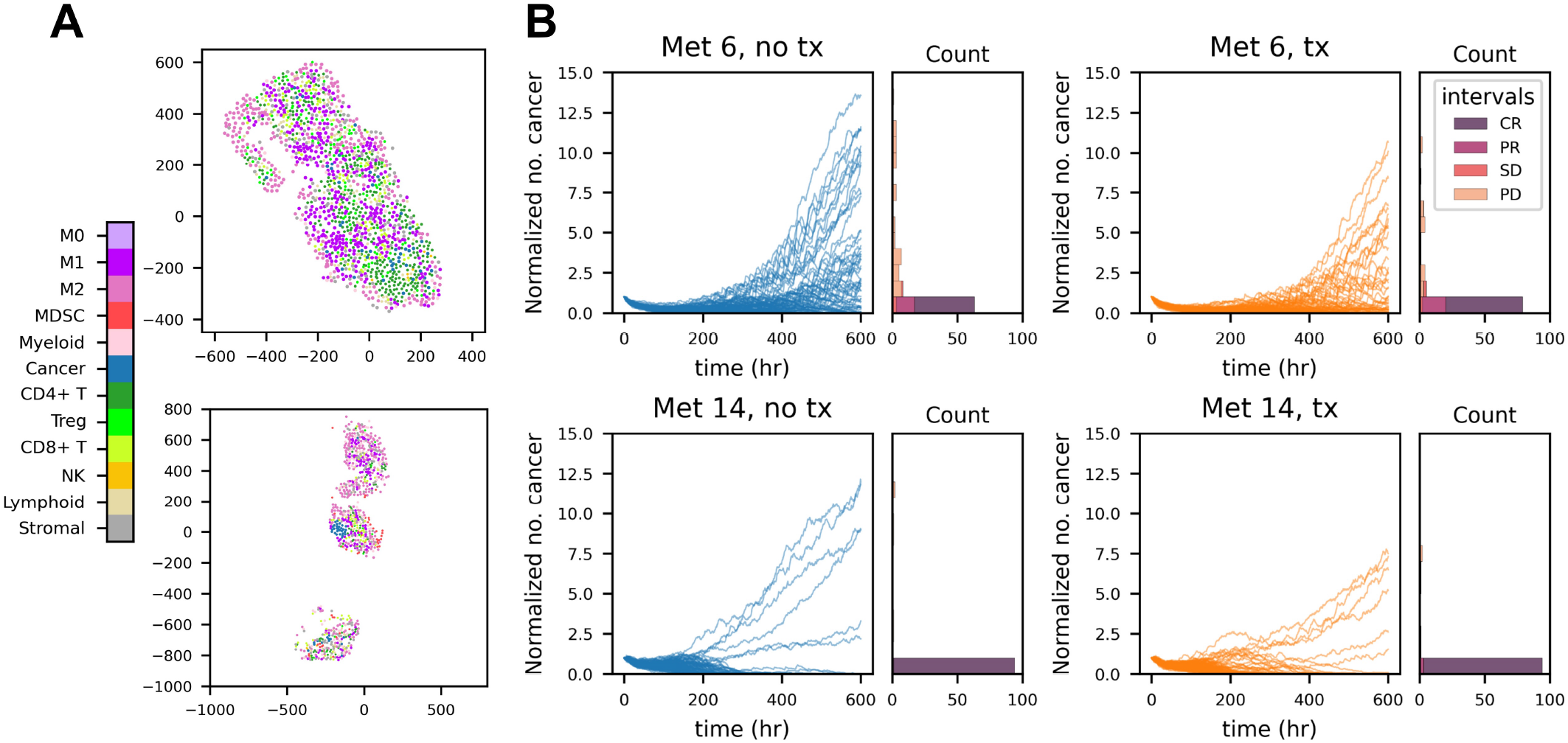
Small, heavily-immune enriched metastases display divergent outcomes. **A.** Initial spatial organization for Met 6 (upper panel) and Met 14 (lower panel). **B.** Cancer growth trajectories for Met 6 and Met 14 (with and without treatment), with the distribution of end states classified by irRECIST-like criteria. One hundred replicates are shown for each metastasis/treatment combination.

**Supplementary Figure S4:**
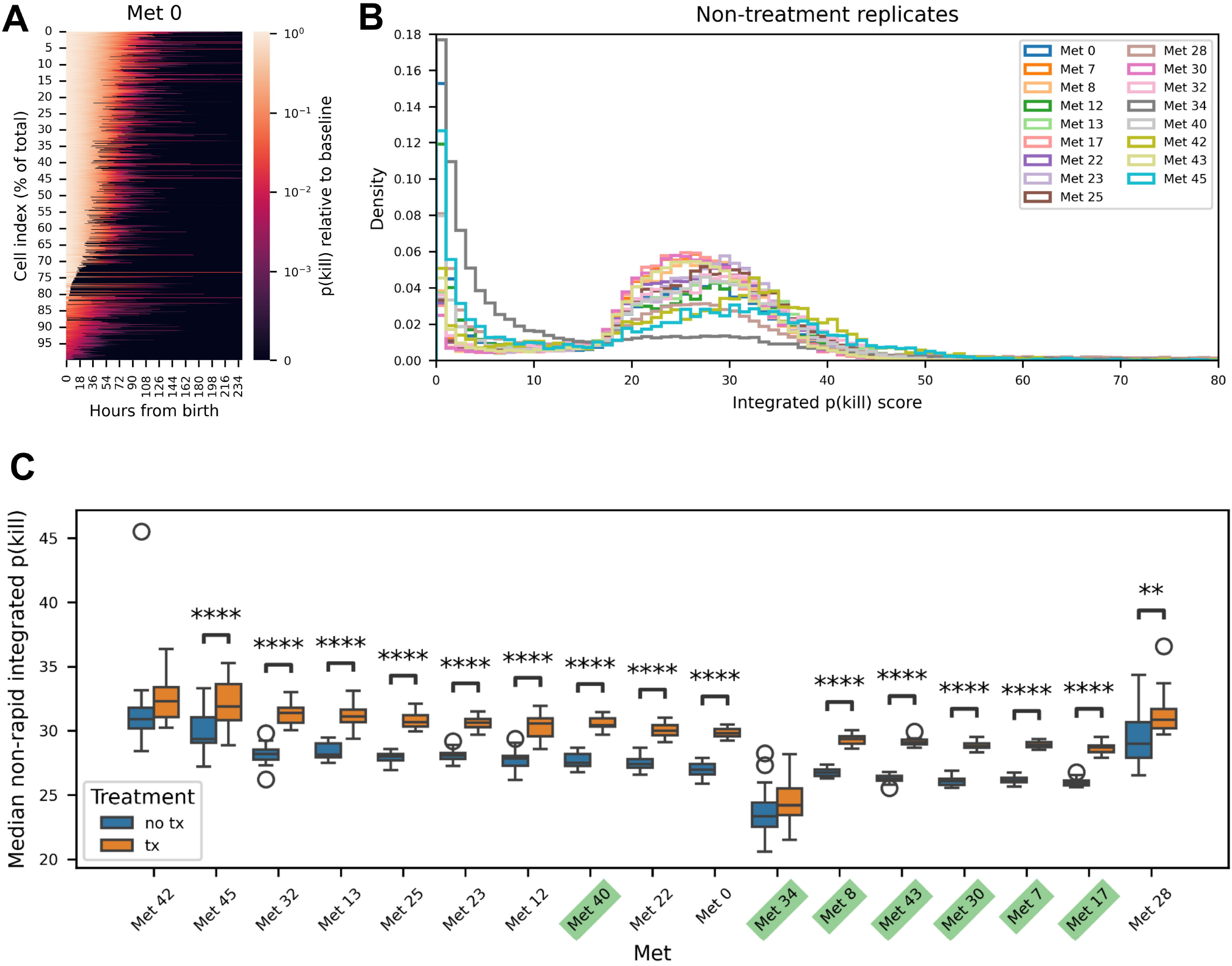
Quantifying bulk CD8+ T cell exhaustion dynamics from simulation illustrates the effect of immune checkpoint inhibition. **A.** An example heatmap showing exhaustion trajectories for the probability of killing for CD8+ T cells in a single replicate (Met 0, no treatment), with lighter coloring indicating higher activity. Cells present at time zero are excluded. **B.** Distribution of CD8+ T cell activity scores obtained by integrating the *p*(*kill*) exhaustion trajectory for no-treatment simulations of metastases. All replicates have been combined to generate these plots. **C.** Boxplots showing that the median integrated *p*(*kill*) score for CD8+ T cells with treatment is higher than for those without treatment across all metastases except for Met 28 and Met 42. Cells with integrated scores below 5 are excluded from this analysis, since they are “rapidly exhausting.” Asterisks denote statistical significance measured by a two-sided student’s *t*-test: * p *≤* 0.05, ** p *≤* 0.01, *** p *≤* 0.001, **** p *≤* 0.0001.

**Supplementary Figure S5:**
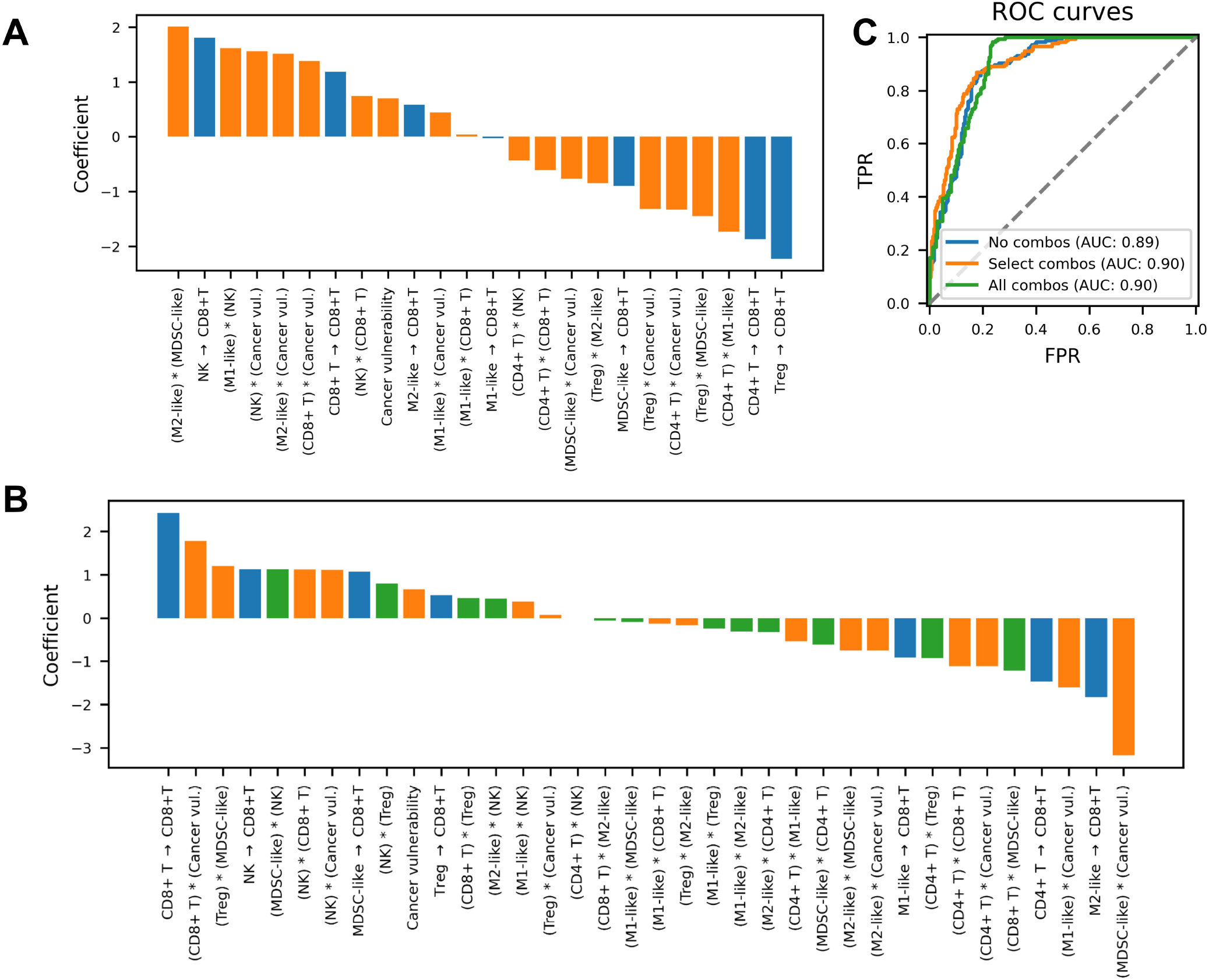
Logistic regression on interaction ratio scores alone performs similarly to logistic regressions that include multiplicative pairwise combinations of interaction ratios. **A.** Coefficients for a logistic regression model incorporating select multiplicative pairwise combinations of interaction ratios and/or cancer vulnerability. Bars in orange represent input features that are not included in the original model described in Fig. 5C termed Regression Model 1, which only contains the seven individual IR scores. **B.** Coefficients for a logistic regression incorporating all multiplicative pairwise combinations of interaction ratios and/or cancer vulnerability, termed Regression Model 2. Bars in orange represent those pairwise combinations included in Regression Model 2 described in **panel A** but not in Regression Model 1 (Fig. 5C), and those in green represent those pairwise combinations only present in this model, termed Regression Model 3. **C.** Receiver operating characteristic curves for all three logistic regression models when trained on the complete data set, showing minimal improvement when adding in select feature combinations and slight improvement when adding in all feature combinations.

**Supplementary Figure S6:**
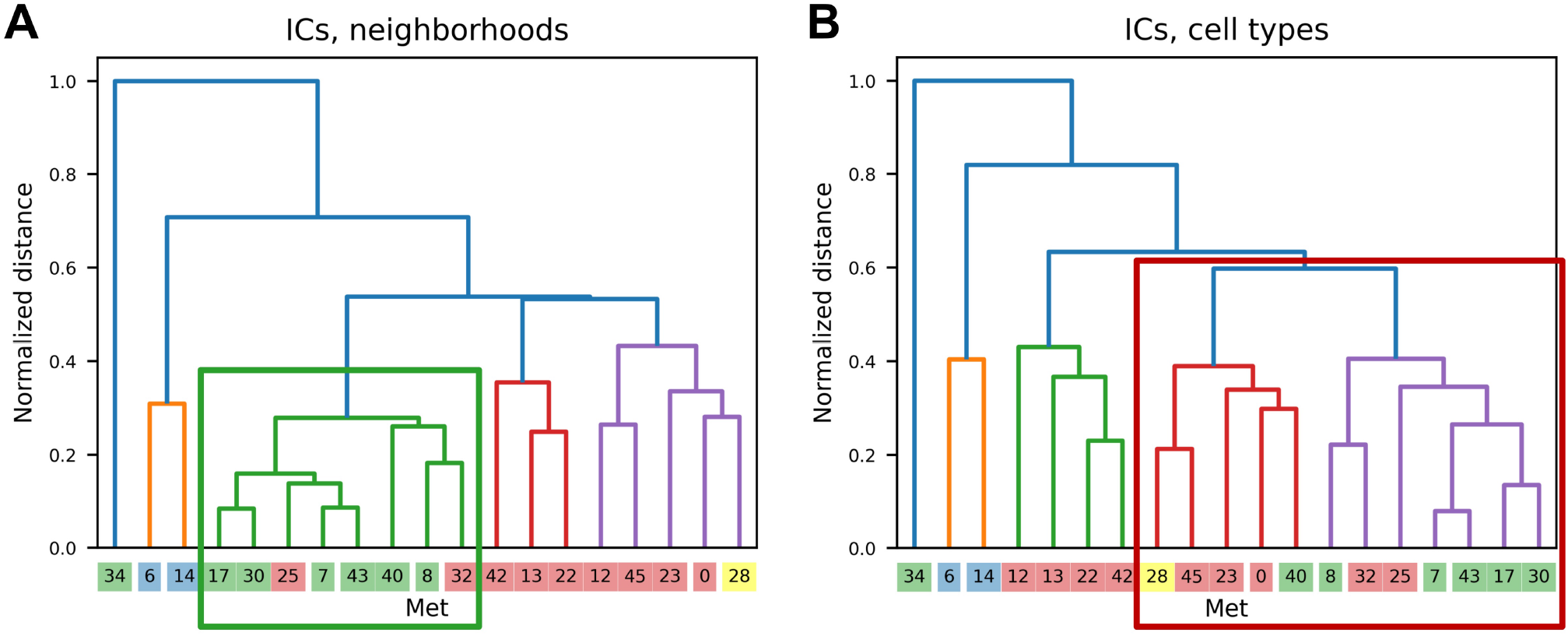
Metastases affected by treatment are better-clustered by initial neighborhood composition compared to initial cell types. **A.** Dendrogram showing hierarchical clustering via initial neighborhood composition based on spatial analysis. The green box denotes the lowest branch containing all metastases affected by treatment except for Met 34. **B.** Dendrogram showing hierarchical clustering via initial cell type composition. The red box denotes the lowest branch containing all metastases affected by treatment except for Met 34.

**Supplementary Figure S7:**
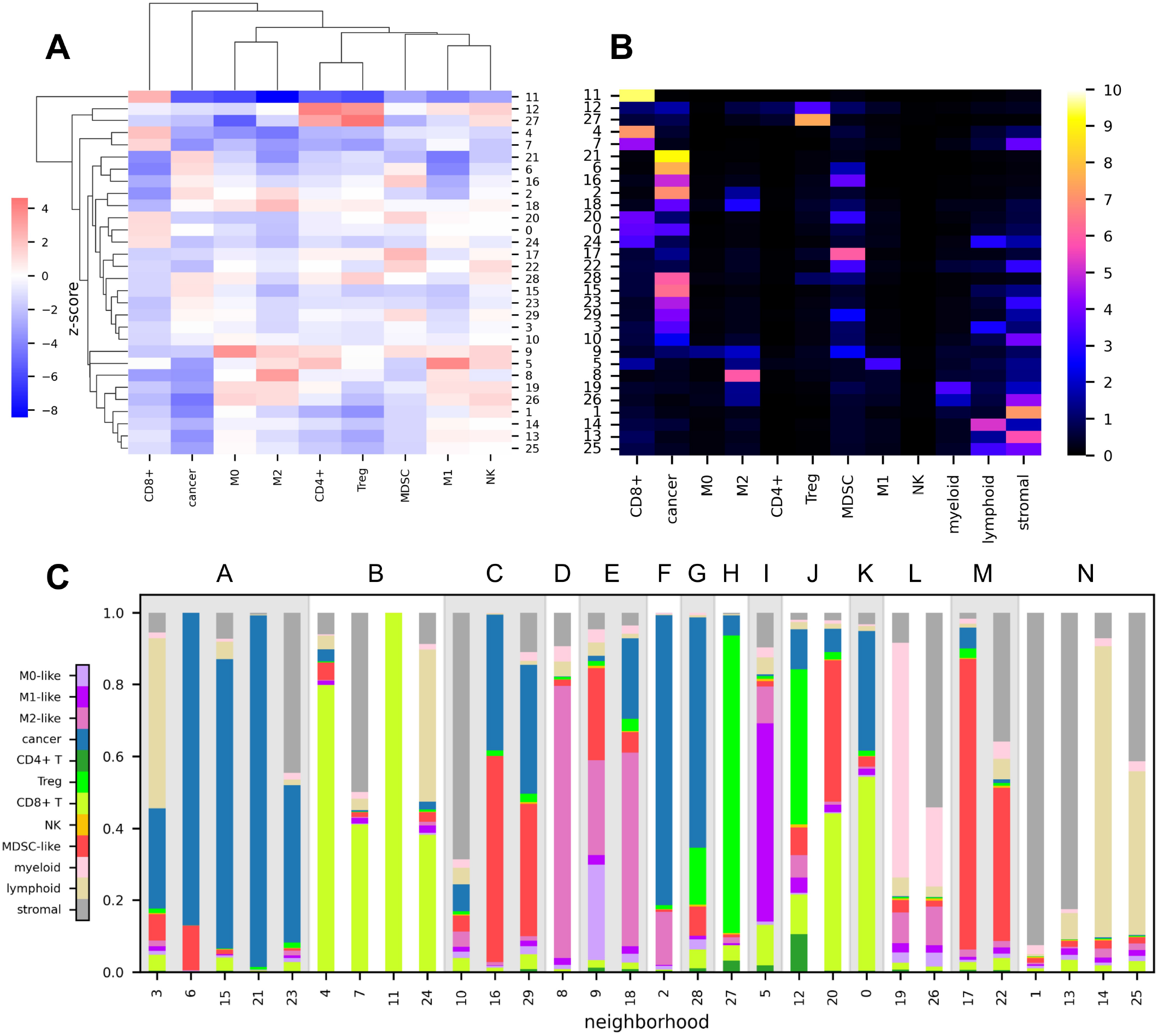
Cellular neighborhood definitions based on the first 48 hours of spatial simulation data in no-treatment replicates. **A.** Heatmap showing the enrichment (Z-score) of different cell types for each cluster (defined by its centroid) relative to the average for all cells in the data set. **B.** Heatmap showing the absolute location of each cluster centroid. **C.** The cell type composition of each neighborhood labeled “A” through “N”. Some neighborhoods are defined based on multiple numbered clusters shown in panels A and B.

