## Supplementary Methods for "Computational modeling of breast cancer metastases predicts response to dual checkpoint inhibition and reveals spatially-informed treatments"

### An agent-based model of tumor-immune interactions in HER2+ breast-to-lung metastases

This document specifies the biological background, scientific justifications, structure, and computational implementation of an agent-based model of lung metastases in a murine model of HER2+ breast cancer that spontaneously seeds metastases to the lungs, liver, and other organs.

### 1 Previous work and summary of the current model

Previously, our group has built agent-based models to investigate the spatiotemporal evolution of tumors in response to various cues in the TME, including the interactions of T cells, macrophages, and cancer cells [1]; and tumor adaptation in response to changes in nutrient, metabolite, and VEGF concentrations, in addition to perturbation under anti-angiogenic therapy [2]. Additionally, we have explored how T cell phenotypes are altered in the TME by coupling an ABM to a Boolean model that predicts CD8<sup>+</sup> T cell states, and analyzing interactions with an expanded immune cell population (CD8<sup>+</sup> T cells, CD4<sup>+</sup> T cells, and macrophages) [3].

In the present model, we expand our work in examining T cell exhaustion in the TME while simultaneously expanding the number of immune cell behaviors we can represent. We integrate two new cell types, natural killer (NK) cells and myeloid derived suppressor cells (MDSCs). Additionally, we focus on two molecular pathways (PD-1 and CTLA-4, see Sections 4 and 5 for more details) that contribute to exhausted phenotypes, and explicitly model how exhaustion changes the behavior of cytotoxic cells. We further model how two immunotherapies that target these pathways, anti-PD-1 and anti-CTLA-4, affect these exhaustion dynamics, in order to understand how cell-cell interactions can contribute to immunotherapy response.

In this document, we provide a complete description of the model. This includes a recapitulation of details carried over from [3], in addition to new additions to the model.

#### 2 Cell types

This agent-based model accounts for six distinct immune cell types, many of which have multiple phenotypes that allow for a diverse array of cell behaviors to be captured (reasoning for why these cell types are included in the model is in Section 6). Generally, immune cell behaviors are classified as “pro-tumor,” which promote suppression of CD8<sup>+</sup> T and NK cells’ cytotoxic capacities, and “anti-tumor,” which promote these cells cytotoxic capacities. Additionally, the model incorporates three “generalized” cell types that allow us to maintain the structural organization of metastases imaged using mIHC where we have low confidence in the identity of the cell based on marker staining.

The behavior of cell agents in the model can be summarized through certain parameters, and many of these behaviors are shared between cells. Key parameters for each cell type are listed in tables in their respective sections, with references for their values, excluding exhaustion parameters for CD8<sup>+</sup> T and NK cells, which are described in Section 5. In this section, we detail these parameters and all cell-intrinsic properties (i.e., those that occur independently of cell-cell interactions). Additionally, we reference processes that are influenced by cell cell interactions, though these are detailed in Section 4.

Additionally, a number of cell parameters (used primarily in the force calculations detailed in Section 3.1) are shared and listed below in Table 1.

| Parameter | Value | Refs. |
| --- | --- | --- |
| Spring constant ($\mu$) | 3 N $\mu\text{m}^{-1}$ | [4] |
| Drag coefficient ($\eta$) | 1 N h$^{-1}$ $\mu\text{m}^{-1}$ | [2] |
| Influence threshold ($x_{\text{th}}$) | 40 $\mu\text{m}$ | Arb. |
| Max overlap | 0.25 | Arb. |
| Attractive force decay constant ($k_c$) | 10 | [2] |

Table 1: Shared parameters for cell behavior

#### 2.1 Macrophages

Macrophages ($M\Phi$) are immune cells in the myeloid lineage that assist in controlling the immune response. In the model, we represent three classes of macrophages: naive macrophages (M0), pro-inflammatory macrophages (M1), and anti-inflammatory macrophages (M2). $M\Phi$s initialized from mIHC data can either be M1 or M2. $M\Phi$s that are recruited to the simulation (see Section 2.8.4) are initialized as M0. Different environmental cues drive the conversion of M0 cells into the M1 or M2 types (a process known as “polarization”). Additionally, M1 and M2 macrophages can interconvert between these polarized states in response to environmental cues. The details for this differentiation process are specified in Section 4.2.

| Parameter | Value | Refs. |
| --- | --- | --- |
| Migration bias | 0.1 | Arb. |
| Migration speed | 72 ($\mu\text{m h}^{-1}$) | [5] |
| M1 transition | 0.4 | Exp. data |
| M2 transition | 0.2 | Exp. data |
| Diameter | 20 ($\mu\text{m}$) | [6] |
| Lifespan | 3 (d) | [3] |
| Recruitment rate | 0.1 (cells $\text{births}_{\text{cancer}}^{-1} \text{ h}^{-1}$) | Exp. data |

Table 2: Parameters for macrophage behavior

#### 2.2 Cancer cells

Cancer cells represent the bulk of the non-immune cells present in our model, and have an expansive ability to proliferate regardless of environmental constraints. Cancer cells proliferate on a clock determined by their internal cell cycle length, which is set either during initialization (from mIHC data, based on the mean and standard deviation in Table 3), or they inherit the cell cycle length from their mother cell following proliferation. The internal cell cycle clock advances regardless of compression effects or the presence of an immune synapse.

| Parameter | Value | Refs. |
| --- | --- | --- |
| Migration speed | 30 ($\mu\text{m h}^{-1}$) | [7] |
| Diameter | 20 ($\mu\text{m}$) | [8] |
| Lifespan | 3.5 (d) | Arb. |
| Cell cycle length (mean) | 30 (h) | Arb. |
| Cell cycle length (std) | 2 (h) | Arb. |

Table 3: Parameters for cancer cell behavior

#### 2.3 CD4<sup>+</sup> T cells

CD4<sup>+</sup> T cells are a class of immune cells in the lymphoid lineage that have a supportive or regulatory role in the immune response. In our model, we have two subclasses of these cells: a CD4<sup>+</sup> helper T cell population (Th), which falls into our “anti-tumor” category and supports the cytotoxic activity of CD8<sup>+</sup> T and NK cells, and a CD4<sup>+</sup> FOXP3<sup>+</sup> T regulatory cell (Treg) that suppresses immune activity and thus falls into our “pro-tumor” category. After initialization, all recruited CD4<sup>+</sup> cells enter the simulation as Th cells, and can *terminally* differentiate into Treg cells in response to certain environmental cues.

| Parameter | Value | Refs. |
| --- | --- | --- |
| Migration bias | 0.2 | Arb. |
| Migration speed | 360 ($\mu\text{m h}^{-1}$) | [9] |
| Differentiation (Th → Treg) | 0.3 | Arb. |
| Diameter | 10 ($\mu\text{m}$) | [8] |
| Lifespan | 3 (d) | [8] |
| Recruitment rate | 0.1 (cells $\text{births}_{\text{cancer}}^{-1} \text{h}^{-1}$) | Exp. data |

Table 4: Parameters for CD4<sup>+</sup> T cell behavior

#### 2.4 CD8<sup>+</sup> T cells

CD8<sup>+</sup> T cells are immune cells in the lymphoid lineage that can target and kill cancer cells (via direct contact and formation of an immune synapse). These cells can proliferate in the same manner as cancer cells (i.e., by using a cell cycle clock), but the cell cycle can be arrested under two conditions. First, if the cell is compressed (see Section 3.1), the cell cycle will not advance. Additionally, if the cell has not made contact with an antigen-presenting cell (APC; either a cancer cell or an M1 macrophage) within a certain time window (see Table 5), then the cell cycle will not advance, and the cell will be unable to proliferate regardless the status of the internal clock. Additionally, the cell cycle length can be increased via the exhaustion process detailed in Section 5, and is re-calculated at each time step. Since we are modeling advanced metastases, we assume that CD8<sup>+</sup> T cells enter simulation in their activated state, and thus can immediately begin killing cancer cells.

| Parameter | Value | Refs. |
| --- | --- | --- |
| Migration bias | 0.2 | Arb. |
| Migration speed | 360 ($\mu\text{m h}^{-1}$) | [5] |
| Diameter | 10 ($\mu\text{m}$) | [8] |
| Lifespan | 14 (d) | [10] |
| Killing probability | 0.1 | Arb. |
| Cell cycle length | 4.5 (h) | [11] |
| Time between antigen presentations | 9 (h) | Arb. |
| Recruitment rate | 0.1 ($\text{cells deaths}_{\text{cancer}}^{-1} \text{ h}^{-1}$) | Exp. data |

Table 5: Parameters for CD8<sup>+</sup> T cell behavior

#### 2.5 Natural killer (NK) cells

Natural killer (NK) cells are immune cells in the lymphoid lineage that can target and kill cancer cells (among others). Unlike CD8<sup>+</sup> T cells, these cells are not proliferative.

| Parameter | Value | Refs. |
| --- | --- | --- |
| Migration bias | 0.125 | Arb. |
| Migration speed | 312 ($\mu\text{m h}^{-1}$) | [12] |
| Diameter | 7 ($\mu\text{m}$) | [13] |
| Lifespan | 14 (d) | [14] |
| Killing probability | 0.1 | Arb. |
| Recruitment rate | 0.1 ($\text{cells deaths}_{\text{cancer}}^{-1} \text{ h}^{-1}$) | Exp. data |

Table 6: Parameters for NK cell behavior

#### 2.6 Myeloid-derived suppressor cells (MDSCs)

Myeloid-derived suppressor cells are immune cells in the myeloid lineage.

| Parameter | Value | Refs. |
| --- | --- | --- |
| Migration bias | 0.1 | Arb. |
| Migration speed | 600 ($\mu\text{m h}^{-1}$) | [15] |
| Diameter | 10 ($\mu\text{m}$) | [16] |
| Lifespan | 2 (d) | [17] |
| Recruitment rate | 0.1 ($\text{cells deaths}_{\text{cancer}}^{-1} \text{ h}^{-1}$) | Exp. data |

Table 7: Parameters for MDSC cell behavior

#### 2.7 Generalized cells

There are three types of generalized cells, which are defined on mIHC data as detailed in Section 6.1. Two of these types are immune cells: the generalized lymphoid cell (which inherits properties from the CD8+ T cell definition) and the generalized myeloid cell (which inherits properties from the MΦ definition). The third type represents a generalized stromal cell. Since we have a low confidence in the identity of these cells, they do not participate in cell-cell interactions and only have a structural function (e.g., they allow the metastasis to maintain its shape during the initial steps of simulation). None of the cell types can proliferate, and the immune-lineage general cells (lymphoid and myeloid) cells die. However, in accordance with other models that span a similar simulation window (i.e., less than 30 days), we do not provide a pathway for stromal cells to die (see references in Table 8); their population remains fixed for the entire simulation.

| Parameter | Generalized Type | Value | Refs. |
| --- | --- | --- | --- |
| Radius | Stromal | 20 ($\mu\text{m}$) | Based on cancer |
| Radius | Lymphoid | 10 ($\mu\text{m}$) | Based on CD8+ T |
| Radius | Myeloid | 20 ($\mu\text{m}$) | Based on MΦ |
| Lifespan | Stromal | ∞ | [18–20] |
| Lifespan | Lymphoid | 14 (d) | Based on CD8+ T |
| Lifespan | Myeloid | 3 (d) | Based on MΦ |
| Migration speed | All | 0 ($\mu\text{m h}^{-1}$) | Manually set |

Table 8: Parameters for generalized cell behavior

#### 2.8 Shared cell processes

##### 2.8.1 Cell death (non-killing)

All cells (excluding generalized stromal cells) can die independently of contact killing events. Cell death occurs as a time-independent process, where a cell has some probability $P$ of dying at each simulation time step defined by the cell’s lifespan parameter (for CD8+ cells, this probability changes as the cell becomes exhausted) and the length of the time step.

When a cell dies, it is removed from the simulation at the *conclusion* of the time step in which it dies. However, once its internal state has been changed to “dead,” the cell can no longer participate in internal processes or cell-cell interaction events.

##### 2.8.2 Proliferation

Both cancer cells and CD8+ T cells can proliferate, and do so using a clock-based mechanism. When the clock exceeds the current cell cycle length, the cell proliferates and resets the cell cycle clock. Upon proliferating, a daughter cell is spawned in a random direction. This direction is produced by a normal vector with $x$ and $y$ coordinates sampled from $\mathcal{N}(0, 1)$, scaled to a distance one cell radius away from the mother cell.

Daughter cells inherit specific mutable properties from their mother cells. Cancer cells inherit PD-L1 expression level and cell cycle length. CD8+ T cells inherit PD-1 expression level, killing probability, migration speed, division probability, and death probability. Thus, CD8+ cells pass on their exhaustion phenotype to daughter cells.

##### 2.8.3 Migration

All cells (excluding generalized cells) migrate within the TME. Additionally, immune cells employ chemotaxis to preferentially migrate toward cancer cells. Thus, we model the migration process as a random walk biased toward the *nearest* cancer neighbor. However, if there are no cancer cells sufficiently close to the immune cell (within 100 $\mu\text{m}$), the cell will not migrate in a biased fashion. This behavior is similar to a cancer cell, which has a bias parameter set to zero. The step size of the random walk is the cell’s speed scaled to the simulation time step length. Thus, for a random unit vector $\vec{v}$ and the unit vector $\vec{u}$ pointing toward the nearest cancer neighbor, the migration vector $\vec{w}$ can be calculated using the bias parameter, $\eta_{bias}$:

$$\vec{w} = \eta_{bias}\vec{u} + (1 - \eta_{bias})\vec{v} \quad (1)$$

The vector $\vec{w}$ is then normalized and multiplied by the cell’s speed to determine the change in position required by the migration process.

##### 2.8.4 Recruitment

Immune cells are recruited into the simulation based on either the number of cancer births ($\text{CD8}^+$ T and NK cells) or the number of cancer deaths (M0, $\text{CD4}^+$ T, and MDSCs) that occurred at the previous time step:

$$n_{\text{recruit}}(t + \Delta t) = \lfloor k_r * n_{\text{cancer births or deaths}}(t) \rfloor$$

When a cell is recruited, it chooses a random location within a rectangle that surrounds the tumor. If the random location is not within a certain distance (100 $\mu\text{m}$) of a cancer cell, the location is rejected and the cell chooses a new random location to spawn, repeating this process until it is within the threshold distance to a cancer cell. The derivation of the scaling parameters $k_r$ for different cell types are described in Section A.2.

#### 3 Physics

##### 3.1 Intercellular forces

Our agent-based model employs the overlapping spheres (OS) method, which represents each cell using two quantities: its center (a coordinate in 2D space) and its radius [21]. Thus, our cells can theoretically occupy any position on the 2D coordinate plane, unlike in grid-based approaches, where cell locations are more restrictive. Using the OS method allows us to integrate more realistic physical processes into our model, especially the physical forces that cells exert on one another. If the vector $\vec{x}_{ij}$ points from the center of cell $i$ to the center of cell $j$, then the force on cell $i$ due to cell $j$ can be determined:

$$\vec{F}_{i,j}(t) = \begin{cases} \mu(r_i + r_j)\hat{x}_{ij} \log_{10}\left(1 + \frac{\|\vec{x}_{ij}\| - (r_i + r_j)}{r_i + r_j}\right) & \text{for } \|\vec{x}_{ij}\| < (r_i + r_j) \\ 0 & \text{for } \|\vec{x}_{ij}\| \geq (r_i + r_j) \end{cases} \quad (2)$$

Additionally, there are cohesive forces between cancer cells, which are added on top of the repulsive forces defined in Eq. 2:

$$\vec{F}_{i,j}(t) = \begin{cases} \mu(|\vec{x}_{ij}| - (r_i + r_j))\hat{x}_{ij} \exp(-k_c \frac{|\vec{x}_{ij}| - (r_i + r_j)}{r_i + r_j}) & \text{for } |\vec{x}_{ij}| \leq x_{\max} \\ 0 & \text{for } |\vec{x}_{ij}| > x_{\max} \end{cases} \quad (3)$$

In the above equations, the value $\mu$ is the spring constant (shared between cells), $r_i$ and $r_j$ are the radii of cells $i$ and $j$ respectively, $k_c$ is the decay constant of the attractive force between cancer cells (as given in Table 3), and $x_{\max}$ is the maximum interaction range. In our model, these forces are used to move cells (in addition to the migration process). This process is solved numerically in Eq. 4, where $\eta$ is the drag coefficient and $\Delta t$ is the simulation step size, which is equal to 0.005 h. The set $\mathcal{N}_i(t)$ is the set of cells within the maximum interaction distance $x_{\max}$ of cell $i$. Here, we refer to the sum of forces on cell $i$ due to cell $j$ as $\vec{F}_{i,j,\text{tot}}(t)$ to indicate that the force vector may consist of attractive, repulsive, and synapse forces (for the latter, see Section 4.1.1).

$$\vec{x}_i(t + \Delta t) = \vec{x}_i(t) + \frac{\Delta t}{\eta} \sum_{j \in \mathcal{N}_i(t)} \vec{F}_{i,j,\text{tot}}(t) \quad (4)$$

##### 3.2 Diffusible factors and cellular influence

Cells within the TME can communicate with each other via diffusible factors, such as cytokines, and affect the local concentration of important metabolic molecules such as glucose and oxygen. These concentrations can be computed discretely by numerically solving systems of partial differential equations, as our group has done in [1] previously. However, as the number of cell types in the model increases along with the number of diffusible factors needed to accurately model cell-cell interactions, it becomes computationally expensive to solve for these diffusible fields. Thus, we employ the method our group previously in [3], where the lumped effect of a cell's secreted diffusibles are modeled using an exponentially-decaying "influence" function (this formulation allows for improved biological fidelity over distance-thresholding methods). We retain the assumption that diffusion occurs more rapidly than cellular processes (i.e., timescale of seconds versus hours), which means that the diffusible factors secreted by a cell will generally "follow" the cell's spatial trajectory and do not persist for long periods of time.

If $|\vec{x}_{ij}|$ is the center-center distance between two cells $i$ and $j$, then the influence of cell $j$ on cell $i$ can be represented by Eq. 5.

$$D_{ij}(t) = \exp(-\lambda|\vec{x}_{ij}|) \quad (5)$$

where $\lambda$ is defined for a distance threshold, $x_{\text{th}}$, and the probability that an effect happens at this distance, $P_{\text{th}}$:

$$\lambda = -\log_2(P_{\text{th}}) \frac{\ln(2)}{x_{\text{th}}}$$

Note, then, that we may interpret the value $D_{ij}(t)$ to represent the probability of an event occurring on cell $i$ due to the influence of cell $j$.

Since the diffusible factors that cells secrete vary by cell type, we calculate values for cell influence by cell type. Let $\mathcal{N}_{i,T}(t)$ denote the set of cells of type $T$ within the neighborhood of cell $i$ at time $t$. Then, the total influence of $T$ on cell $i$ can be given by Eq. 6.

$$D_{i,T}(t) = 1 - \prod_{j \in \mathcal{N}_{i,T}(t)} (1 - D_{ij}(t)) \quad (6)$$

This value represents the probability that an event occurs on cell $i$ due to all cells of type $T$, and the equation falls out of the fact that we consider these events to be independent of one another (the product represents the probability that *no event occurs*, and thus the probability that *any* event occurs is one minus this product).

We can use a similar logic to represent the combined influences of multiple cell types, shown in Eq. 7.

$$D_{i,T_1+T_2}(t) = 1 - (1 - D_{i,T_1})(1 - D_{i,T_2}) \quad (7)$$

#### 4 Cell-cell interactions

Cell-cell interactions (CCIs) form the core of our model. There are two major classes of interactions in this model: direct interactions, which require physical contact between two cells to take place, and indirect interactions, which are mediated by diffusible factors represented by cell influences. Additionally, we describe a special subset of direct interactions that are mediated by immune synapses. These CCIs directly module a number of important cell processes, including macrophage and $CD4^+$ T cell differentiation, and impact $CD8^+$ T and NK cell exhaustion, as detailed in Section 5. Further, they control the expression of PD-1 and its ligand PD-L1 on various cell types, which facilitate the PD-1-PD-L1 exhaustion program.

##### 4.1 Direct interactions

Direct interactions can occur between cells when they overlap; that is, when:

$$||\vec{x}_{ij}|| < r_i + r_j \quad (8)$$

Direct interactions only occur between specific sets of cells. In this section, when a cell of type $A$ “undergoes” interactions with a cell of type $B$, it means that the direct contact with $B$ has caused some effect on $A$, but $B$ is not necessarily affected. We also call $A$ the “target” cell and $B$ the “effector” cell, as in Table 9 below, which summarizes these processes (but excludes immune synapse-mediated killing which involve cancer, $CD8^+$ T, and NK cells, which are detailed in Section 4.1.1).

MDSCs, $CD4^+$ T cells, $M\Phi$, and generalized cells are not targets for direct interactions, though the first three types can be effectors for direct interactions.

| Target | Effector | Effect |
| --- | --- | --- |
| CD8<sup>+</sup> T | cancer, M1 | antigen contact activation |
| CD8<sup>+</sup> T | cancer, M2, Treg, MDSC | PD-1/PD-L1 mediated exhaustion process |
| NK | cancer, M2, Treg, MDSC | PD-1/PD-L1 mediated exhaustion process |

Table 9: Summary of direct interactions

###### 4.1.1 Immune synapses

Immune synapses (also called immunological synapses) are a major feature of many immune cell interactions, and describe a physical synapse that forms between two immune cells or an immune cell with some other target cell. For example, cytotoxic T cells form two major types of immune synapses: lytic synapses, which are used to kill target cells, and stimulatory synapses, which can lead to T cell activation. Following the formation of a lytic synapse, target cells quickly die as lytic granules are quickly pushed into the target cell after only a few minutes of contact [22], prompting apoptotic signaling within the target which can lead to cell death after approximately 30 minutes [23]. Lytic synapses are typically thought to degrade soon after target death (within 1-2 hours in [22]), but stimulatory synapses typically have a longer duration (e.g., 2 to 12 hours) [23]. Additionally, researchers have observed CD8<sup>+</sup> T cells forming multiple lytic synapses and polarizing lytic granules toward multiple targets without competition between the synapses [22].

Additionally, NK cells can form immune synapses with their targets. These synapses are structurally similar to CTL-target synapses, undergo a similar formation process, and possess a similar killing efficacy, though their duration is shorter [24].

Depending on immune context, the presence of PD-1 in a CTL-target immune synapse can lead to either destabilization (e.g., autoimmune contexts) or stabilization (e.g., antiviral response) of the synapse complex, and in this latter case, stabilization can lead to an apparent “paralysis” of the T cell since it cannot disengage to kill other targets [25]. Additionally, the presence of PD-1 in lytic synapses can impair the actin remodeling needed to release lytic granules, meaning that CTL cytotoxic capacity can be diminished by PD-1/PDL-1 interactions [26]. Other work, however, has shown that α-PD-1 antibodies can exclude PD-1 from the immune synapse (in addition to blocking PD-1/PDL-1 signaling), and thus lead to improved T cell activation [27].

In our model, we allow CD8<sup>+</sup> T and NK cells to synapse with their target cells only (i.e., we model lytic synapses, not stimulatory synapses for T cells or inhibitory synapses for NK cells). Because these the model evaluates major events at a timestep of $\Delta t = 1$ h, we set the duration of all immune synapses to one hour, and this value does not change with exhaustion or PD-1 expression. Additionally, since the immune synapse physically connects the cells together, we introduce an additional force that pulls the cell toward each other. This force is modeled as a spring force, where the spring has a resting length slightly smaller than the sum of the two cells’ radii (i.e., this spring enforces some small overlap between the cells), as in Eq. 9.

$$F_{i,j}^{\vec{x}}(t) = -\hat{x}_{ij}k_{\text{syn}}(l_{\text{rest}} - ||\vec{x}_{ij}||) \quad (9)$$

Additionally, the resting length of the spring is defined:

$$l_{\text{rest}} = r_i + r_j - l_{\text{overlap}}$$

In the model, the parameter values for the spring constant $k_{\text{syn}}$ is set to be stronger ($10 \text{ N}\mu\text{m}^{-1}$) than that of the spring constant used in Eq. 2 ($\mu = 3 \text{ N}\mu\text{m}^{-1}$), so the synapse force dominates the repulsive force caused by the overlap. Note that this spring force must be added to the total forces used to resolve the change in position caused by cell forces, as shown in Eq. 4.

Additionally, we limit $\text{CD8}^+$ T and NK cells to forming a maximum of two immune synapses, as observed in [22]. When the hour-long lifespan of the synapse elapses, we delete the connection and the cell is free to form a new synapse with the same or different cell. Since these cells employ the synapse to deliver lytic granules to targets, $\text{CD8}^+$ T and NK cells *must* be synapsed with a target in order to kill it. At every major time step, cytotoxic cells attempt to kill their synapsed targets and succeed based on the cell's intrinsic killing probability.

#### 4.2 Indirect interactions

Cells undergo indirect interactions due to diffusible molecules secreted by other cells within a neighborhood, as described by Eqs. 5 - 7. These indirect interactions control things such as PD-1 and PD-L1 expression, exhaustion effects (discussed in Section 5, marked (EX) in the table), and differentiation. These interactions and their effects are summarized in Table 10.

| Target | Effector | Effect |
| --- | --- | --- |
| $\text{CD8}^+$ T | cancer, M2, Treg, MDSC | PD-1 expression |
| $\text{CD8}^+$ T | M1, $\text{CD4}^+$ T | (EX) ↑ killing probability, migration speed |
| $\text{CD8}^+$ T | M2, Treg, MDSC | (EX) ↓ killing probability, migration speed |
| $\text{CD8}^+$ T | $\text{CD4}^+$ T, NK | (EX) ↑ proliferation rate |
| $\text{CD8}^+$ T | M2, MDSC | (EX) ↓ proliferation rate |
| NK | cancer, M2, Treg, MDSC | PD-1 expression |
| NK | M1, $\text{CD4}^+$ T | (EX) ↑ killing probability, migration speed |
| NK | M2, Treg, MDSC | (EX) ↓ killing probability, migration speed |
| Cancer | $\text{CD8}^+$ T, $\text{CD4}^+$ T, NK | PD-L1 expression |
| M2 | $\text{CD8}^+$ T, $\text{CD4}^+$ T, NK | PD-L1 expression |
| Treg | $\text{CD8}^+$ T, $\text{CD4}^+$ T, NK | PD-L1 expression |
| MDSC | $\text{CD8}^+$ T, $\text{CD4}^+$ T, NK | PD-L1 expression |
| All $\text{M}\Phi$ | $\text{CD8}^+$ T, $\text{CD4}^+$ T, NK | M1 polarization |
| All $\text{M}\Phi$ | cancer, M2, Treg, MDSC | M2 polarization |
| $\text{CD4}^+$ T | cancer, M2, MDSC | Differentiation into Treg |

Table 10: Summary of indirect interactions

##### 4.2.1 PD-1 and PD-L1 expression

The induction of PD-1 expression on $CD8^+$ T and NK cells is controlled by the cancer cells, M2 macrophages, Tregs, and MDSCs. Thus, we can express the positive influences on PD-1 expression following the form of Eq. 7:

$$D_+ = 1 - (1 - D_{\text{cancer}})(1 - D_{\text{M2}})(1 - D_{\text{Treg}})(1 - D_{\text{MDSC}})$$

We then compare $D_+$ to some threshold probability ($P_{\text{th, PD-1}}$), and if it exceeds this value, we increase the PD-1 expression inherent to the cell (unless it reaches a saturation value). However, if the threshold is not exceeded, which may occur if there are insufficient cells of the required types in the neighborhood then the PD-1 decays according to zero-order kinetics (where $v_{\text{PD-1}}$ is the rate of PD-1 induction and $d_{\text{PD-1}}$ is the constant rate of PD-1 decay):

$$\text{PD-1}(t + \Delta t) = \begin{cases} \text{PD-1}(t) + (D_+ - P_{\text{th, PD-1}})v_{\text{PD-1}}\Delta t & \text{for } D_+ \geq P_{\text{th, PD-1}} \\ \text{PD-1}(t) - d_{\text{PD-1}}\Delta t & \text{for } D_+ < P_{\text{th, PD-1}} \end{cases}$$

The induction of PD-L1 on cancer cells, M2 macrophages, Tregs, and MDSCs is controlled in a similar manner by $CD4^+$ T $CD8^+$ T and NK cells. In this case, we express the positive influences of PD-L1 expression as:

$$D_+ = 1 - (1 - D_{\text{Th}})(1 - D_{\text{CD8}})(1 - D_{\text{NK}})$$

Then, we follow the same form to determine the change in PD-L1 expression inherent to the cell, with the same zero-order decay process where $v_{\text{PD-L1}}$ is the rate of PD-L1 induction and $d_{\text{PD-L1}}$ is the constant rate of PD-L1 decay:

$$\text{PD-L1}(t + \Delta t) = \begin{cases} \text{PD-L1}(t) + (D_+ - P_{\text{th, PD-L1}})v_{\text{PD-L1}}\Delta t & \text{for } D_+ \geq P_{\text{th, PD-L1}} \\ \text{PD-L1}(t) - d_{\text{PD-L1}}\Delta t & \text{for } D_+ < P_{\text{th, PD-L1}} \end{cases}$$

##### 4.2.2 Differentiation processes

Both $M\Phi$s and $CD4^+$ T cells can undergo a differentiation process where they transform from one set of behaviors (which define a cell “subtype”) to another. This contrasts, for example, with the exhaustion process that affects $CD8^+$ T and NK cells, where the affected properties exist on a continuum (see Section 5), rather than as discrete subtypes.

The two $CD4^+$ subtypes in our model are helper cells (denoted $CD4^+$ T) and T regulatory cells (Treg). Additionally, $CD4^+$ T cells can undergo an *irreversible* transition to become T regulatory cells. The probability that a $CD4^+$ T cell differentiates into a Treg is governed by the influence of cancer cells, M2 macrophages, and MDSCs, and a scaling constant ($p_0$) which is the probability differentiation occurs when at least one of the influencing cells has $D = 1$:

$$p_{\text{diff}} = p_0 D_{\text{diff}},$$

$$D_{\text{diff}} = 1 - (1 - D_{\text{cancer}})(1 - D_{\text{M2}})(1 - D_{\text{MDSC}})$$

In our model, MΦs have three states: the naïve M0 state, the anti-tumor M1 state, and the pro-tumor M2 state. M0 macrophages can differentiate into either of the two polarized subtypes (M1 or M2), but once polarized, macrophages cannot reenter the naïve M0 state. Thus, M1 macrophages can only differentiate into M2 macrophages and vice versa. Mathematically, macrophage differentiation is governed by the *relative probabilities* of transition events, where $p_i$ is the relative probability that a macrophage will transition into the $M_i$ state (from any state, so this also could represent the probability that the macrophage *does not* change its polarization). Like with $CD4^+$ T cells, the relative probabilities for M1 and M2 transition are given by scaled influence functions, while the probability $p_0$ is nonzero (and equals 1) only for M0 macrophages.

$$\begin{aligned} p_0 &= \begin{cases} 1 & \text{if } M0 \\ 0 & \text{if } M1, M2 \end{cases} \\ p_1 &= k_{M1}(1 - (1 - D_{Th})(1 - D_{CD8})(1 - D_{NK})) \\ p_2 &= k_{M2}(1 - (1 - D_{cancer})(1 - D_{M2})(1 - D_{Treg})(1 - D_{MDSC})) \end{aligned}$$

We then normalize the relative probabilities by dividing by their sum ($p_0 + p_1 + p_2$) to obtain the normalized transition probabilities such that $\hat{p}_0 + \hat{p}_1 + \hat{p}_2 = 1$.

#### 5 Exhaustion dynamics

In the tumor microenvironment, chronic stimulation by antigens can lead to a phenomenon called “exhaustion,” where T effector cells lose their cytotoxic capacity, upregulate inhibitory receptors, and undergo a host of other physical and behavioral changes [28]. Additional research has revealed that NK cells, which can kill without antigen presentation, also experience this phenomenon, mediated by many of the same pathways (especially through inhibitory receptors) [29]. Noticeable upregulation of exhaustion markers have been observed to occur on T cells within hours of their arrival to the TME [30], and we evaluate the cell processes and cell-cell interactions that govern exhaustion at every major timestep (1 h).

In our model, the behaviors of $CD8^+$ T and NK cells are governed by five major parameters: killing probability, migration speed, migration bias, life span, and cell cycle length. Generally, a property $p$ are updated according to Eq. 10 where the function $D(t)$ represents some set of cell influences *or* the effect of PD-1/PD-L1 interactions, and the scaling parameter $k_d$ is the single source exhaustion decay constant (described in Section A.1). NK cells do not proliferate in our model, so we do not track a cell cycle length for them.

$$p(t + \Delta t) = (1 - D(t)(1 - k_d))p(t) \quad (10)$$

In the case where cells are exhausted from a single source (e.g. $D(t) = 1$), the above equation results in exponential decay for each property (we track the *inverse* of the cell cycle length and lifespan properties). The decay constants for each property are given in Table 12, and their derivations from the literature sources cited are in section A.1. We use a slightly modified version of the equation to track the lifespan, since we model death as a probability:

$$p_{\text{death}}(t + \Delta t) = (1 + D(t)(k_d^{-1} - 1))p_{\text{death}}(t)$$

Our model employs two major immunosuppressive pathways that contribute to exhaustion: CTLA-4-mediated suppression (which only affects CD8<sup>+</sup> T cells), and PD-1/PD-L1-mediated suppression (which affects CD8<sup>+</sup> T and NK cells).

#### 5.1 CTLA-4-mediated exhaustion

Cytotoxic T-lymphocyte-associated antigen 4 (CTLA-4) is an immune checkpoint protein expressed on T regulatory cells that participates in cytotoxic T cell inhibition. Unlike the PD-1/PD-L1 axis, where PD-L1-bearing cells attenuate effector cell function via direct contact, the CTLA-4 pathway impacts cytotoxic T cells indirectly by preventing their activation by antigen-presenting cells (APCs) like dendritic cells. This occurs because CTLA-4 can bind to CD80/86 molecules present on APCs, which alternatively bind to the costimulatory molecule CD28 on effector cells [31, 32]. The CD28-CD80/86 interaction is necessary for effector cell activation; thus, competition for CD80/86 on APCs between CTLA-4 and CD28 leads to diminished effector cell function. Indeed, CTLA-4 has a higher binding affinity for CD80/86 relative to CD28.

Because our model does not explicitly contain APCs that participate in T cell activation, we model the effect of CTLA-4 and its associated immune checkpoint inhibitor indirectly. Specifically, we allow the influence of T regulatory cells to negatively affect the proliferation probability of CD8<sup>+</sup> T cells. The proliferation of these cells are affected negatively and positively by different immune cells as described below:

$$\begin{aligned} D_+ &= 1 - (1 - D_{\text{Th}})(1 - D_{\text{NK}}) \\ D_- &= 1 - (1 - D_{\text{M2}})(1 - D_{\text{MDSC}}) \end{aligned}$$

The effect of the anti-CTLA-4 drug is modeled by a Hill equation, where we call $\eta$ the “effect” of the drug. Because activated CD8<sup>+</sup> T cells may have upregulated CTLA-4 expression, we assume that cells with a cell cycle length shorter than baseline are more sensitive to the drug, and multiply the Hill effect by an arbitrary coefficient $k_{\text{act}}$ equal to 1.1 (the coefficient is one when the cell cycle length is equal or greater than baseline).

$$\eta = k_{\text{act}} \left( 1 + \left( \frac{\text{EC50}}{[\alpha\text{-CTLA-4}]} \right)^n \right)^{-1}$$

Then, from the form of Eq. 10, we have:

$$\begin{aligned} D(t) &= D_- - D_+(1 - D_{\text{Treg}}(1 - \eta)) \\ p(t + \Delta t) &= (1 - D(t)(1 - k_d))p(t) \end{aligned}$$

Note that the form of the $D_+$ term above is *different* than the multiplicative influence effect described by Eq. 6; instead, the Treg influence term *inhibits* the pro-proliferative interactions represented by $D_+$ (e.g., when the Treg influence is one and $\eta = 0$ because there is no drug, the total $D_+$ term is zero). In the case where the drug fully occupies all the CTLA-4 site on the Treg, we have $\eta = 1$, and this inhibitory effect is negated.

Additionally, we assume that severely exhausted cells are resistant to the α-CTLA-4 drug, and thus have $\eta = 0$. We define severely exhausted as having a killing probability, migration speed, and lifespan one-quarter that of baseline.

#### 5.2 PD-1/PD-L1-mediated exhaustion

The expression of programmed cell death protein 1 (PD-1) is regarded as a hallmark of T cell exhaustion, both marking and leading to decreases in effector cell function. Its ligand, PD-L1, can be expressed on cells like macrophages and MDSCs, and binding between PD-1 and PD-L1 impairs signaling at the T-cell receptor [32]. Additionally, many cancer cells can acquire PD-L1 expression via environmental stimuli (for example, due to interferon gamma secretion by effector cells) [33]. The PD-1/PD-L1 pathway can also affect NK cells, leading to the attenuation of their effector function [29].

The description of how environmental factors influence the induction of PD-1 and PD-L1 in different cell types is above in Section 4.2.1. In order for the PD-1/PD-L1 exhaustion process to occur, cell must be in direct contact, e.g. following Eq. 8. We then assume a simple 1-to-1 binding between process PD-1 and PD-L1 that occurs until one of the molecules has been saturated. In our model, we represent the concentrations of these surface molecules on a unitless scale from zero to five, since the calculations we perform are agnostic to the actual expression values.

In the presence of α-PD-1, the amount of PD-1 that its ligand can bind to is reduced. We assume that the drug and the surface receptor quickly reach equilibrium, and thus the fraction of PD-1 unoccupied by the drug can be derived from standard kinetics.

$$\% \text{ PD-1}_{\text{available}} = 1 - \frac{[\alpha\text{-PD-1}]}{[\alpha\text{-PD-1}] + K_D}$$

We can then describe the effect of PD-1/PD-L1 interactions on the property values in question, by defining $D(t)$ and using it in Eq. 10. Here, we scale the effect of PD-1/PD-L1 binding such that if half of the maximum number of PD-1 site are bound to their ligand, the “scale factor”

$$D(t) = \min([\text{PD-1}_{\text{available}}], [\text{PD-L1}]) \frac{[\text{PD-1}_{\text{max}}]}{2}$$

As with the α-CTLA-4 drug, we assume that severely exhausted cells (same definition) are resistant to the drug.

#### 5.3 Indirect exhaustion

While PD-1/PD-L1 mediated exhaustion require direct contact (e.g., receptor-ligand binding), diffusible signals can also promote (or mitigate) exhaustion in $\text{CD8}^+$ T and NK cells.

$$\begin{aligned} D_+ &= 1 - (1 - D_{\text{M1}})(1 - D_{\text{Th}}) \\ D_- &= 1 - (1 - D_{\text{M2}})(1 - D_{\text{Treg}})(1 - D_{\text{MDSC}}) \end{aligned}$$

In both cytotoxic cell types, we the update three properties through indirect exhaustive processes – killing probability, migration speed, and migration bias – according to the balance of positive and negative (anti- and pro-exhaustive) influences.

$$p(t + \Delta t) = p(t)k_d^{(D_+ - D_-)}$$

#### 6 Integrating experimental data

We initialize our simulations using cell type and location data obtained from multiplexed immunohistochemistry (mIHC) imaging of mouse lung metastases. Imaging is carried out using the following markers, which allow resolution of the different non-generalized cell types described in this model: HER2, CD3, CD8, FoxP3 GR1, MHCII, CD206, NCR1. Following segmentation, all cells were assigned binary expression labels in addition to the fluorescence intensity values measured for different cell areas (e.g., nucleus, membrane, cytoplasm). Initial cell type labels were assigned by mapping known binary expression patterns to their corresponding cell types. However, this led to a large number of cells being assigned to an “unknown” group.

##### 6.1 Random forest labeling of unidentified cells

Cells assigned an unknown cell type via their protein expression were assigned a cell type using a random forest (RF) model. The random forest model was trained using the cells successfully labeled using protein expression. The input features for the random forest were the fluorescence values from mIHC. Unlabeled cells were assigned either a defined cell type or a generalized cell type based on the class probabilities predicted by the RF model (see **Sup. Fig. S1** in the **Main text**). Briefly, if the highest class probability is large, or sufficiently large relative to other classes, then the cell is assigned into that cell type. However, if this is not the case, then we examine the relative (summed) probabilities of the immune lineages to determine whether or not a cell should be assigned into the generalized types of “myeloid” or “lymphoid,” or if it should be assigned to the non-immune “stromal” class.

##### 6.2 Scaling

The radii of the various cells in our model are taken from the literature, rather than being approximated from mIHC images. Often, however, there are initial states that result in large amounts of overlap between cells when using these literature values. Because key cancer and CD8<sup>+</sup> T cell features are changed when these cells are compressed, this can lead to significant simulation artifacts during early time points that are biologically unrealistic. In order to mitigate these simulation artifacts, we scale mIHC coordinates prior to simulation such that *on average*, cells are not compressed at $t = 0$. Scaling the coordinates directly, rather than allowing the simulation to determine a spatial equilibrium, is preferable because it maintains the observed geometries of the metastasis.

To calculate the scale factor, we compute the estimated actual density, $\rho_a$ and estimated required density, $\rho_t$ for the metastasis. To calculate $\rho_a$, we first estimate the area of the metastasis by computing the alpha-shape of the cell centers with a sufficient $r$ parameter to enclose the entire met with a single contour. We consider the alpha-shape to be the boundary of the metastasis, and its enclosed area is the area of the metastasis. The density $\rho_a$ is then simply the number of cells divided by the enclosed area.

To calculate $\rho_t$, we consider the total area that would be covered by all cells in the met given our predefined radii. Because circles cannot fully cover the plane, however, we then scale each area by a constant $k = \frac{\sqrt{12}}{\pi}$ to account for this fact (the constant is derived from the percentage of the plane that hexagonally-packed circles occupy). Thus, for a metastasis with $n$ cells, we can write:

$$\rho_t = \frac{n}{k \sum_{i=1}^n (\pi r_i^2)} \quad (11)$$

Finally, we obtain the scale factor $s$ by which we multiply each coordinate:

$$s = \sqrt{\frac{\rho_a}{\rho_t}} \quad (12)$$

##### 6.3 Initialization from images

The processed, post-scaling data is saved as .csv files that contain cell ID, x and y coordinate positions, and the cell type. This data is passed directly into the model, which generates initial conditions using the coordinate positions and cell type. Since we do not acquire property-specific information (beyond cell type) from mIHC imaging, we must generate information like PD-1 and PD-L1 expression when models are initialized, as specified in Table 11.

| Property | Cell types |
| --- | --- |
| PD-1 expression level | CD8<sup>+</sup> T, NK |
| PD-L1 expression level | cancer, Treg, MDSC, M2 |
| Cell cycle length | cancer, CD8<sup>+</sup> T |
| Cell cycle position | cancer, CD8<sup>+</sup> T |
| Cell age | CD8<sup>+</sup> T, NK |
| Migration speed | CD8<sup>+</sup> T, NK |
| Lifespan | CD8<sup>+</sup> T, NK |
| Killing probability | CD8<sup>+</sup> T, NK |

Table 11: Cell properties randomly generated during initialization.

For non-cytotoxic (e.g., not CD8<sup>+</sup> T or NK) cells, their properties are generated as follows:

• A cell’s PD-1 or PD-L1 expression level is determined by uniformly sampling the interval between 0 and the maximum expression level;
• A **cancer** cell’s cell cycle length is sampled from a normal distribution;
• A **cancer** cell’s cell cycle position is determined by uniformly sampling the interval between 0 and the cell’s assigned cell cycle length.

CD8<sup>+</sup> T and NK cells also have their PD-1 expression levels set by uniformly sampling the interval between 0 and the maximum expression level. However, exhaustion-related parameters (migration speed, lifespan, killing probability, cell cycle length) are determined by the randomly-generated age of the cell, to model the temporal effect of exhaustion.

While exhaustion is both temporally and spatially path-dependent, we make a number of simplifying assumptions in order to generate the expected age distribution. First, we assume that cells experience a constant amount of exhaustion prior to simulation time zero (this is the same assumption we make when deriving exhaustion-related parameters). From prior simulations, we have observed that cells exhaust (on average) at a rate 1.5 times higher than would be expected from a single exhaustion source. Thus, the probability of death (the hazard function) will approximately be the exponential function:

$$p_{death}(t) = \alpha \exp(1.5(k_d - 1)t)$$

Here, $\alpha$ denotes the death probability for an unexhausted cell, and $k_d$ is the inverse of the lifespan decay parameter. This means that distribution of lifespans that we should observe will follow a Gompertz distribution of the form:

$$\alpha \exp\left(\frac{\alpha}{1.5 * k_d} + 1.5 * k_d t - \frac{\alpha}{1.5 * k_d} \exp(1.5 * k_d t)\right)$$

Accordingly, we can generate a random lifespan, $l$, using the quantile function of the Gompertz distribution and a random variable $X$ which is uniformly sampled from the interval $[0, 1]$:

$$l = \frac{1}{1.5 * k_d} \ln\left(1 - \frac{1.5 * k_d}{\alpha} * \ln(1 - X)\right)$$

We then assume that we can recover the expected distribution of ages by randomly sampling the lifespan distribution and converting a random lifespan into a random age by uniformly sampling the age from the interval $[0, l]$. Once we have determined the age of the cell, we can then set each of the exhaustion parameters using the different $k_d$ values associated with each property:

$$p = p_{base} \exp(-a * (1 - k_d))$$

#### 7 Model implementation

The model described in the previous sections has been implemented using C++, and the most recent version can be accessed via GitHub. In short, we have created a program which takes mIHC data and a number of parameters as inputs, and outputs results from simulation (as described in Section 7.5 below). In this section, we give a brief overview of how the model is structured in C++ and important computational processes (e.g., RNG generation). We do not present the code in detail, but note that we use Python to import parameters into the model, so running the code requires both C++ and Python. Additionally, running the code in parallel requires OpenMP, and the program is currently configured to be built and deployed in a Linux-based system.

#### 7.1 Cell agents are objects

As agent-based models describe the evolution of a number of discrete agents over time, they are well suited for object-oriented programming. Each cell in the simulation is represented by an object, which is an instance of its cell type class. Each cell type class is derived from a shared class, which contains shared functions for actions like migration, PD-1 and PD-L1 expression, and contact inhibition. A number of functions which are common to multiple cell types, but have different behaviors for each, are also defined here. However, their definitions are empty and must be overridden by the child class. Note that differentiation processes do not change the class of the cell object; instead, differentiation changes an internal state variable.

#### 7.2 Stochastic events

In the agent based model, all events that proceed “probabilistically” are evaluated stochastically through the seeded random number generator described below.

Consider an event that occurs with a probability $P$. We then generate a random number (RNG) on a uniform distribution from zero to one (e.g. $U(0, 1)$). There are then two cases:

$$\begin{aligned} \text{RNG} \leq P &\rightarrow \text{event occurs} \\ \text{RNG} > P &\rightarrow \text{event does not occur} \end{aligned}$$

When we have “relative probabilities” ($P_1, P_2, \dots, P_n$), which occurs when an event has more than two outcomes (e.g., macrophage polarization), we generate a random number (RNG) on a uniform distribution from zero to one and then evaluate:

$$\begin{aligned} 0 \leq \text{RNG} < P_1 &\rightarrow \text{event 1 occurs} \\ P_1 \leq \text{RNG} < P_1 + P_2 &\rightarrow \text{event 2 occurs} \\ &\vdots \\ P_1 + \dots + P_{n-1} \leq \text{RNG} \leq 1 &\rightarrow \text{event } n \text{ occurs} \end{aligned}$$

Normal distributions are used to assign certain cancer cell properties upon initialization (i.e., cell cycle length), to choose locations for cell proliferation, and to determine the random component for cell migration.

In addition to informing probabilistic events as described above, uniform distributions are used to distribute cell birth times, PD-1 expression levels, and PD-L1 expression levels when initializing cells from imaging. Additionally, we use uniform distributions to randomly assign exhausted phenotypes to CD8<sup>+</sup> T and NK cells when initializing from imaging.

#### 7.3 Random number generation

In order to make our results reproducible, our agent based model contains a random number generator which can be seeded through a user-defined base seed. Generally, when we show a distribution containing $n$ runs, the seeds we use are the integers on the interval $[0, n - 1]$. In addition to the main RNG instance used in sequential regions of the program, we also create a number of temporary RNG instances when the program runs in parallel across multiple CPU cores. The base seed $H_{\text{temp}}$ for a temporary instance is deterministically calculated using the simulation timestep, the unique cell ID (since parallel distribution is done on a per-agent basis), and the process ID (preset for different model processes):

$$\begin{aligned} H_1 &= \text{base seed} \\ H_2 &= H_1 \oplus (\text{time step} + h_1 + (H_1 \ll 6) + (H_1 \gg 2)) \\ H_3 &= H_2 \oplus (\text{cell ID} + h_2 + (H_2 \ll 6) + (H_2 \gg 2)) \\ H_{\text{temp}} &= H_3 \oplus (\text{process ID} + h_3 + (H_3 \ll 6) + (H_3 \gg 2)) \end{aligned}$$

Above, $h_1, h_2$, and $h_3$ are hash constants (0x9e3779b9, 0x9e3779b3, and 0x9e3779b1, respectively), $\oplus$ represents the XOR function, and ≪ and ≫ are the left and right bit-shifts respectively.

The RNG object can generate random numbers (either integers or doubles) according to uniform, normal, and Poisson distributions. We use a Mersenne Twister 19937 generator to get numbers, seeded using either the base seed or the temporary base seed.

#### 7.4 Simulation loop

After set-up functions, the program enters the main simulation loop. For each time step, of length $\Delta t_1 = 1$ h (the “$\Delta t_1$ time step”), the following high-level processes are evaluated sequentially (regions of code where the individual agent processes are parallelized are marked with $^\dagger$, otherwise subprocesses within each enumerated item occurs sequentially):

1. Drug dynamics are approximated,
2. Immune cells are recruited based on the previous step’s cancer births and deaths,
3. Neighboring cells are determined$^\dagger$,
4. Influences and indirect interactions are calculated$^\dagger$,
5. Direct interactions are calculated$^\dagger$,
6. Differentiation processes occur$^\dagger$,
7. Internal cell properties are updated,
8. The high-resolution (with $\Delta t_2$) migration loop occurs $\Delta t_1/\Delta t_2$ times,
9. Cells die due to age,
10. Cells proliferate (if possible),
11. Dead cells are removed from simulation (their objects are recorded then deleted),
12. The order of the cell list is shuffled,
13. Simulation outputs are recorded, and
14. The simulation determines whether or not to terminate.

The high-resolution migration loop, which has a time step of length $\Delta t_2 = \frac{1}{200}$ h (the “$\Delta t_2$ time step”), contains the following sequence of steps, with individual agent processes fully parallelized within each step:

1. Cells migrate,

2. Neighboring cells are determined,
3. Intercellular forces are calculated,
4. Intercellular forces are resolved,
5. Neighboring cells are re-determined,
6. Immune synapses are formed and/or updated, and
7. Cell compression is evaluated.

The simulation will terminate if 1) the cancer population reaches zero (e.g., the metastasis is eliminated), or 2) the simulation reaches the maximum time step, which is set to $t = 25$ d.

###### 7.4.1 Avoiding data races

During certain parallelized regions of code (see above), interactions between cells can result in internal properties being changed *while other cells are attempting to access those properties*. Since we dynamically allocate threads in these regions (and sometimes vary the number of cores we use to run simulations), these “data races” may result in untraceable stochastic effects that prevent reproducibility and have unintended effects on biology. Thus, we do not immediately update cell properties when they are changed through direct or indirect interactions with other cells, and instead store them in an “update” member. After the interaction processes are completed, we then replace cell properties with their corresponding “update” member, thus avoiding data race conditions. Likewise, when we calculate (biased) random walks for cells, we do not update the location of cells until all cells have determined their next position.

#### 7.5 Outputs

As the program runs, it will generate various outputs that record the simulation’s progress. Additionally, we record the parameters passed into the simulation.

At the conclusion of each $\Delta t_1$ time step, the simulation records a snapshot of all cells present (cells which have died during this timestep are removed prior to data capture, so are not recorded). Two files store this information: `cells.csv` and `influences.csv`. At the conclusion of a simulation which has run for $n$ $\Delta t_1$ time steps, there should be $n$ different `cells.csv` and `influences.csv` files storing the spatial and phenotypic information of all cell agents for each time step:

• In `cells.csv`, rows correspond to different cell agents and we record the following seven values: unique cell ID, cell type, X coordinate, Y coordinate, radius, cell state, mother unique cell ID. For CD8<sup>+</sup> T and NK cells, we also store six more values: PD-1 expression level, available PD-1, migration speed, killing probability, death probability, and division probability. The “death probability” recorded is the inverse of the cell’s life span, and the “division probability” is the inverse of the cell’s cell cycle length. Additionally, for the four exhaustion parameters, we do not record their absolute value, but rather their ratio with the base value of the parameter. For all other cells, we record the PD-L1 expression level and the cell cycle length (absolute value), and pad the remaining four columns with -1.

• In `influences.csv`, rows again correspond to different cell agents, and we record the unique cell ID followed by the influences exerted on that cell by other cell types in the following order: M0, M1, M2, cancer, Th, Treg, CD8+, CD8+ (repeated), NK, NK (repeated), MDSC.

Additionally, the program generates four files that store population-level time series data about the simulation:

• The `cancer_death.csv` file stores the number of cancer cells which have died at each $\Delta t_1$ time step due to age, CD8<sup>+</sup> T cell killing, and NK cell killing.
• The `cd8_proliferation.csv` file stores the number of CD8<sup>+</sup> T cells which have proliferated at each $\Delta t_1$ time step.
• The `immune_contact.csv` file stores the total number of cancer cells in contact with an immune cell and immune cells in contact with a cancer cell at each $\Delta t_1$ time step. Note that this is *not* the number of immune synapses that have formed, since “contact” simply requires that the cells are touching or overlap.
• The `populations_TS.csv` file stores the total number of each cell type present in the simulation at each $\Delta t_1$ time step in the following order: M0, M1, M2, cancer, Th, Treg, CD8, NK, MDSC, myeloid, lymphoid, stromal. Additionally, the file stores the computed tumor radius in its last column.

If the α-PD-1 or α-CTLA-4 drug are present in simulation as either monotherapy or combination therapy, the program will generate files that track the concentration of the drug in the simulation environment at each $\Delta t_1$ time step.

It is also possible to recover the *positions* for each cell agent for every $\Delta t_2$ time step, though this is by default disabled since these files tend to be quite large. If this is re-enabled, files labeled by unique cell ID will be generated for each cell and store the coordinate location of the cell’s center. It is currently not possible to recover any other information at a $\Delta t_2$ time step resolution.

### A Parameterization

#### A.1 Exhaustion parameters

As described above, we assume that we can model the effect of exhaustion on important cytotoxic cell parameters as some form of exponential decay. To derive the decay constant, we assume that at each time step the cell experiences the same exhaustive effect. As mentioned above, NK cells do not proliferate in this model, so we do not model the impact of exhaustion on their reproduction rate, or the effect of NK cell proliferation on exhaustion. The values obtained through this parameterization process are summarized below in Table 12.

| Cell type | Property | $k_d$ | Half-life |
| --- | --- | --- | --- |
| CD8<sup>+</sup> T cell | Killing probability | 0.957 | 16 h |
| CD8<sup>+</sup> T cell | (Cell cycle length)<sup>-1</sup> | 0.982 | 39 h |
| CD8<sup>+</sup> T cell | Velocity | 0.979 | 32 h |
| CD8<sup>+</sup> T cell | Life span | 0.9897 | 67 h |
| CD8<sup>+</sup> T cell | Migration bias | 0.979 | 32 h |
| NK cell | Killing probability | 0.984 | 43 h |
| NK cell | Velocity | 0.992 | 32 h |
| NK cell | Life span | 0.9897 | 67 h |
| NK cell | Migration bias | 0.992 | 32 h |

Table 12: Derived exhaustion decay constants for CD8<sup>+</sup> and NK cells

##### A.1.1 T cell cytotoxicity

First, to approximate the decay in CD8<sup>+</sup> T cell cytotoxicity we examine the data presented in [30], which quantifies the number of effector cells bearing a TNF<sup>+</sup> INF-γ<sup>+</sup> phenotype at different time points. We use this percentage as a proxy for the killing probability relative to baseline, since we may interpret this value as a population-level representation of the killing capacity of all effector cells at the time of measurement.

Working from the raw data provided in the supplementary material, we run a regression on the log-transformed data shown in Fig. 4g of [30] (we only consider the effector cells that are transferred into mice bearing lung metastases, e.g. “E→T”), excluding an outlier present in the 48 h data. We assume that this model has a value of one at time zero, which is fixed. This calculation yields a decay constant of 0.957 h<sup>-1</sup>, which corresponds to a half-life of approximately 16 h. Because our model assumes that T cells are appropriately activated before entry into the metastasis (given that these are late-stage mets), we choose this data set over the alternative presented in Fig. 4b of this paper.

##### A.1.2 NK cell cytotoxicity

NK cells also exhaust upon entering the tumor microenvironment, which has been shown to occur quickly, with the level of cell markers associated with exhausted phenotypes correlated with the residence time of the NK cell within the tumor [34]. To quantify the change in cytotoxic capacity of NK cells within our model, we use the data presented in [34], where the authors divide the NK cell population into a pre- and post-48-hour group using photo-conversion and quantify the amount of cells in each age group that land in sc-RNA clusters associated with cell surface markers. The differential gene expression and pathway analyses demonstrate how important genes related to NK cell function change over time, but it is difficult to approximate quantitative values from the data provided. For these experiments, cells are harvested 24, 48, and 72 hours post-photoconversion, which means that the Kaede-red labeled cells may have been present for up to 3 to 5 days, depending on which harvest group they are in.

In the data presented in Fig. 2J from the paper, we can observe the proportion of cells in different groups (green and red photolabeled 24 and 72 hours post photoconversion) expressing specific markers—for example, the exhaustion marker LAG3 which corresponds to restricted production of interferon gamma and reduced degranulation [35, 36]; and the post-activation marker CD69+, which can approximate the number of cells in each compartment that have been activated. The photolabeled cells are grouped through their expression of CD11b<sup>+</sup> (indicating more mature NK cells [37]) and CD49a<sup>+</sup> (indicating “tissue-resident” cells). We will use change in the expression of LAG3 to approximate the decay in cytotoxic capacity in the NK cell population, where the proportion of LAG3<sup>+</sup> cells approximates the percent reduction in killing probability for these cells.

While these groupings do not, of course, persist over time (since cells could gain or lose CD11b<sup>+</sup> and CD49a<sup>+</sup> expression over the course of the experiment), we can approximate the change in LAG3 expression between the green populations and red populations for cell group (CD11b<sup>+</sup>, double-negative, and CD49a<sup>+</sup>). This yields five approximations (we do not use the CD11b<sup>+</sup> Kaede green group since the initial condition is essentially zero) for the exponential decay coefficient, with an average of 0.984, or a half-life of 49 hours.

##### A.1.3 T cell proliferation

Exhaustion is further characterized by reduced proliferative capacity, especially in the terminally exhausted population ($T_{EX}$; this is relative to a more proliferative progenitor-exhausted population, $T_{PEX}$, which is a precursor state to $T_{EX}$). While [30] presents data from a CFSE assay which allows very coarse-grained estimation of how proliferation changes over a 60 h window, their numerical data are not well suited to deriving a decay constant as we did above. Rather, we turn to [38], which also uses a CFSE assay to examine differences in $T_{PEX}$ and $T_{EX}$ subsets in a mouse model of melanoma, and slices these data into one-division wide bins (except for the last bin, which is 7+ divisions; see Fig. 3e in the paper). After using ImageJ to determine the height of each bar in Fig. 3e from the paper, we can estimate the average number of divisions each population experienced over 96 hours, which can then be transformed back into a cell cycle length. From this data, we determined that $T_{PEX}$ cells have a cell cycle length of 16.3 h, while $T_{EX}$ cells have a cell cycle length of 21.7 h.

While we do not have clean time point data as we had from [30] above, we can still attempt to estimate the rate at which the CD8<sup>+</sup> T cell’s proliferative capacity changes. The cells in [38] were harvested significantly later than those in [11]. The data in [30] suggests that there is an initial period following adaptive transfer during which the CD8+ cells do not proliferate (e.g. the first 12 hours), after which they begin to divide with increasing rapidity, even though these cells accumulate markers of exhaustion like PD-1 prior to this interval in other sections of the study (and display exhausted phenotypes for cytotoxic effects). However, we do not know for how long the TEX and TPEX cells have been resident in the TME for, just that they are subsequently cultured in vitro for four days. If we accept that CD8+ T cells have a 14-day lifespan, then we might conclude that those cells have existed in the TME for no longer than 10 days, after which point the cells do not experience the suppressive effects of the TME nor further antigen contact; however, since exhausted T cells have shortened lifespans, it is likely that these cells are younger. Estimates for the cell cycle length of CD8+ T cells following activation (the “expansion phase”) are estimated to be between 6 to 8 hours [11], though faster doubling times have been observed and used in other models (e.g., 4.5 hours) [39, 40].

We first consider the case where the initial doubling time for an effector cell upon entry to the tumor is 4.5 h. We then consider the last time point in [30] (48 to 60 hours) and assume the longest doubling time here, which is around 13.3 h. We then calculate the coefficient necessary for exponential decay to achieve this value at the specified time, which results in a multiplier of 0.9821 hr<sup>-1</sup>. A cell on this trajectory achieves the proliferation-based definition of T<sub>PEX</sub> at 71.3 h (2.97 days) and the definition of T<sub>EX</sub> at 87.1 h (3.63 days), which are both within the 10 day window we determined via [38]; this result gives us the quickest decay for proliferation rate we might expect in the model. We next consider the case where the initial doubling time for an effector cell is 7 h. We now assume a slightly higher doubling time at 60 h, equal to 8.28 h. This yields a coefficient equal to .9946 h<sup>-1</sup>. With these parameters, a cell will reach the T<sub>PEX</sub> proliferation level at 186 hr (7.75 days) and the T<sub>EX</sub> proliferation level at 239.5 hr (9.98 days). This means a cell will reach the T<sub>EX</sub> levels at the latest possible point. While there are other combinations of parameters (initial cell cycle length and exponential decay coefficient), these examples provide bounds on what combinations might lead to the values observed in these studies.

###### A.1.4 T cell apoptosis

Exhausted cells can also undergo early apoptosis, leading to decreased lifespans within the TME. For example, research shows that CD8+ T cells that have exhaustion-associated markers PD-1 and LAG3 have increased active caspase-3, which indicates an activated apoptotic program [41]. Other work shows that exhausted T cells express increased levels of the proapoptotic protein Bim and that the Bcl2/Bim ratio in this exhausted subset is lower than in non-exhausted CD8+ T cells [42].

The data from [41] is suggestive of increasing quantities of active caspase-3 as exhaustion progresses, though we expected exhausted cells to generally be older than non-exhausted cells, since it takes time for T cells to accumulate multiple antigen hits. While there is no actual time-structuring in this data, we can see that the percentage of active caspase-3 rises from 1.2% to 20% when moving from PD1<sup>-</sup> LAG3<sup>-</sup> 4-1BB<sup>-</sup> to PD1<sup>+</sup> LAG3<sup>+</sup> 4-1BB<sup>+</sup> (the CD8<sup>+</sup> T cells are from mice that have borne tumors for 13 days prior to harvesting). When all the cells from a tumor are pooled, the average percent active caspase-3 is 9.17%, which is slightly lower than what is reported for the PD1<sup>+</sup> LAG3<sup>+</sup> group; thus meaning that the PD1<sup>+</sup> LAG3<sup>+</sup> 4-1BB<sup>+</sup> group is over twice as apoptotic relative to the average T cell (which has a phenotype similar to the PD1<sup>+</sup> LAG3<sup>+</sup> group). The data in [42] suggests that cells gaining a different apoptosis-associated marker after exhaustion (Rgs16)

can undergo apoptosis as early as 6 days following Rgs16 expression. Again, none of this data is adequately time structured, but we use it to inform the assumption that a cell experiencing constant exhaustion since its birth should have a lifespan of closer to 6 days. In this case, we choose our scaling coefficient by making it such that for a random death process (evaluated every hour), the probability that a cell with no exhaustion is alive at day 14 is the same as the probability that a cell with a single exhaustion source is alive at day 6. The equation used to obtain this value is:

$$\prod_{n=0}^{336} \left(1 - \frac{1}{336}\right) = \prod_{n=0}^{144} \left(1 - \frac{1}{336(k_d)^n}\right)$$

Solving this equation gives that $k_d = 0.9897$. Because we do not have comparable data for NK cells, we set the NK cell's $k_d$ to this value as well.

##### A.1.5 NK and T cell motility

Changes to CD8+ T cell motility within the TME seems to involve multiple, conflicting processes. For example, researchers have shown that T cell velocities within solid tumors decrease due to mechanical stiffness caused by ECM crosslinking [43]]. Both CTLA-4 and PD-1 have been shown to alter how T cells interact physically with other cells, which may change their motility and apparent velocities, though this relationship is complex. For example, work has shown that exhausted T cells (measured through PD1 expression) have observed velocities up to twice that of non-exhausted T cells [44]. Additionally, surface molecules like CTLA-4 can also increase T cell motility by disrupting binding to dendritic cells, and by overriding the TCR-induced stop signal [45]. Other work, however, has shown that anti-CTLA-4 and anti-PD-1 can increase cell motility, and the authors conclude this may occur because these drugs disrupt weak bindings between T cells and APCs like dendritic cells [45]. We have already integrated immune synapses into our model, which will slow the apparent velocities of cytotoxic cells engaged with their targets. However, since there is conflicting data in the literature, we have arbitrarily set the decay for CD8+ and NK cell velocities to be half that of their cytotoxic capacity (i.e., $(1 - k_{d,\text{velocity}}) = 0.5(1 - k_{d,\text{killing}})$).

Additionally, recent reports have further shown that the T cell exhaustion program impacts not only the velocity at which these cells move, but also leads to down-regulation of genes involved in chemotaxis that may be upregulated following acute antigen stimulation (rather than chronic) [46, 47]. We do not directly model chemotaxis in the model, but the bias parameter in the random walk approximates this process, and thus allows us to control the relative “strength” of chemotaxis as exhaustion proceeds. However, we do not have quantitative data to inform the rate at which we decrease the migration bias in the random walk. Therefore, for both cell types, we set it to the same rate of decay as their respective decays in migration speed.

#### A.2 Recruitment parameters

Recruitment of immune cells is proportional to the number of cancer deaths and/or births at each time step. To determine the rates of recruitment for each immune cell type, we analyze the data from each metastasis and determine the recruitment rate (relative to the number of cancer births and deaths that would occur in that moment) necessary for the ratio of immune cells to cancer cells to remain constant.

Let $C$ denote the number of cancer cells in a given metastasis, and $b_c$ and $d_c$ denote the birth and death rates respectively. Then, we have that:

$$\frac{dC}{dt} = C(b_d - d_c)$$

To maintain the proportion of some other immune cell population, $I$, we then must have:

$$\frac{dI}{dt} = \frac{I}{C} \frac{dC}{dt}$$

We can also express the change in $I$ due to birth, death, and recruitment. The parameters $b_I$ and $d_I$ are the birth and death rates of $I$ respectively, and $r(I)$ evaluates to $b_c$ if the immune recruitment is cancer-birth dependent or $d_c$ if the immune recruitment is cancer-death dependent:

$$\frac{dI}{dt} = k_r * r(I) * C + I(b_I - d_I)$$

We can then solve for $k_r$ by setting the two expressions for $\frac{dI}{dt}$ equal:

$$k_r = \frac{I}{r(I) * C} ((b_c - d_c) - (b_I - d_I))$$

We then solve for these across the 19 mets we are currently analyzing. Because the composition of the mets vary significantly, and there are some mets which possess zero of a specific cell type, we take the geometric mean of the rates with zero-values excluded (see below for the log-scaled distributions; additionally, we plot the time-structured data but do not see any differences between the different harvest times, mostly because the vast majority of the data was from the same harvest timepoint). (Note that we also set the proliferation rate of CD8<sup>+</sup> T cells to zero to represent a strong exhaustive environment; we can adjust this value and get various scaling factors for CD8<sup>+</sup> T cell recruitment).
